# CRootBox: A structural-functional modelling framework for root systems

**DOI:** 10.1101/139980

**Authors:** Andrea Schnepf, Daniel Leitner, Magdalena Landl, Guillaume Lobet, Trung Hieu Mai, Shehan Morandage, Cheng Sheng, Mirjam Zörner, Jan Vanderborght, Harry Vereecken

## Abstract

**Background and Aims:** Root architecture development determines the sites in soil where roots provide input of carbon and take up water and solutes. However, root architecture is difficult to determine experimentally when grown in opaque soil. Thus, root architectural models have been widely used and been further developed into functional-structural models that simulate the fate of water and solutes in the soil-root system. We present a root architectural model, CRootBox, as a flexible framework to model architecture and its interactions with static and dynamic soil environments.

**Methods:** CRootBox is a C++ -based root architecture model with Python binding, so that CRootBox can be included via a shared library into any Python code. Output formats include VTP, DGF, RSML and CSV. We further created a database of published root architectural parameters. The capabilities of CRootBox for the unconfined growth of single root systems, as well as the different parameter sets, are highlighted into a freely available web application.

**Key results:** We demonstrate the use of CRootBox for 5 different cases (1) free growth of individual root systems (2) growth of root systems in containers as a way to mimic experimental setups, (3), field scale simulation, (4) root growth as affected by heterogeneous, static soil conditions, and (5) coupling CRootBox with Soil Physics with Python code to dynamically compute water flow in soil, root water uptake, and water flow inside roots.

**Conclusions:** In conclusion, we present a fast and flexible functional-structural root model which is based on state-of-the-art computational science methods. Its aim is to facilitate modelling of root responses to environmental conditions as well as the impact of root on soil. In the future, we plan to extend this approach to the aboveground part of the plant.

## 1 Introduction

Root architecture development determines the sites in soil where roots provide input of carbon and energy and take up water and solutes. Thus, plant roots strongly interact with their soil environment (Gregory, 2006). However, root architecture is difficult to determine experimentally when grown in opaque soil. Therefore, root architectural models have been widely used for generating root architectures for a large variety of plants and been further developed toward functional-structural models that are able to simulate the fate of water and solutes in the soil-root system.

Root architecture models may be distinguished into three broad levels of complexity. Root depth models (that assume an exponential root length distribution over depth, Raats, 1974), density-based root models (Dupuy *et al*. 2010, Roose *et al*. 2001), and 3D root architectural models that take into account dynamic development of root structure (e.g. Leitner *et al*. 2010a). It was recognised very early that impact of roots on soil processes should be taken into account in soil models. Often this impact was modelled using simple parameterizations of root related processes such as root water uptake (Feddes *et al*. 1978) and plant nutrient uptake (Somma *et al*. 1998). Root architecture models on the other hand were initially used to visualize and analyse the branched structure of root systems which was otherwise not observable in opaque soil (Diggle 1988; Lynch *et al*. 1997; Pagès *et al*. 2004). Over time, “function” was added to those structural root architecture models (e.g. Dunbabin *et al*. 2002), while structural root architectural models have been merged with soil models (e.g. Javaux *et al*. 2008). Both approaches have now been merged to complex functional-structural models that are able to simulate the fate of water and solutes in the soil-root system (Dunbabin et al., 2013, Chimungu and Lynch 2014, Schröder *et al*. 2014), some including rhizosphere gradients around each root segment (Schnepf *et al*. 2012, Schröder *et al*. 2009) or hydraulic and chemical signalling (Huber *et al*. 2014).

Today, applications are needed at a range of spatial scales, requiring information about root systems, including single plant and crop models, field scale models as well as regional and larger scale models such as land surface models. Some of those models suffice with root length density information for the computation of e.g. root water uptake sink terms. Other models solve water, solute and carbon flow inside roots as well. In those cases, the root segment length is an important parameter as it controls the discretisation of the numerical grid on which flow and transport equations are solved, where stability and convergence conditions such as the Courant-Friedrichs-Lewy condition or the von Neumann condition need to be fulfilled. In this work we describe a framework to simulate the response of root architecture to soil environmental properties as well as the influence of roots on soil conditions in a dynamic way. The paper is organized in the following manner.

In the section materials and methods, we present the C++-based root architecture model CRootBox, which fulfils these criteria. It is based on the earlier RootBox code that had been implemented in Matlab (Leitner *et al*. 2010), but now has an object oriented implementation which is more flexible and faster so that field-scale modelling is now feasible.

The key differences of CRootBox with respect to other root architecture models include

- Root segments have a user-defined length
- Results are independent of spatial and temporal resolution
- Easy interface to be coupled with other (e.g. soil) models
- Fast, works from single root to plot scale
- Fast analysis tools included
- Confining containers or obstacles are considered based on signed distance functions

The focus of CRootBox is the simulation of different types of root architecture, and to provide a generic interface for coupling with arbitrary soil/environmental models, e.g., in order to determine the impact of specific root architectures on their functions, e.g. related to drought resistance or nutrient uptake efficiency. To demonstrate this generic interface, we show an example of coupling with a code from the book “Soil Physics with Python”. The coupling to the soil model is realised with Python. The Python binding is realised with the C++ library Boost. Python. Output formats include VTK Polygonal Data format (VTP), which can be visualised in Paraview, a plain text file containing coordinates of root nodes as well as the Root System Markup Language (RSML) format developed by (Lobet et al., 2015) which is now widely used in a number of different image analysis, general root analysis and modelling tools (e.g. Excel, R, R-SWMS, CRootBox) and the Dune Grid Format (DGF) which can be used by the generic partial differential equation solving environments Dune and DuMu^x^ (Flemisch *et al*. 2011).

The structure of CRootBox is outlined in Fig. 1. CRootBox is a dynamic root architecture model that can respond to soil conditions. Static heterogeneous soil conditions can be defined in CRootBox via signed distance functions or via look-up tables. Furthermore, CRootBox provides generic interfaces to interact with external models that simulate e.g. the soil environment or root internal states.

**Fig. 1.**
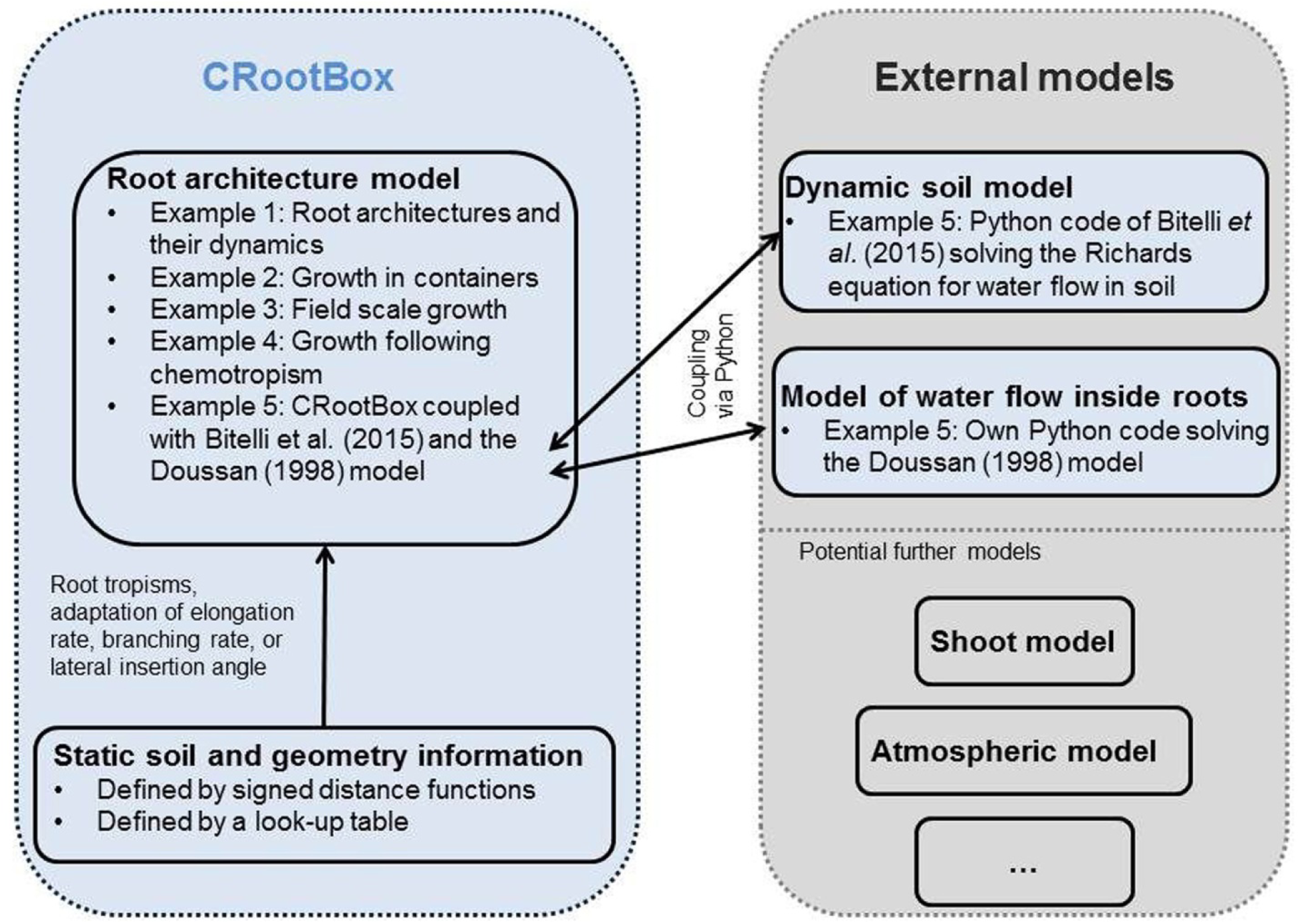
CRootBox is a dynamic root architecture model that can respond to soil conditions. It provides generic interfaces to interact with external models that simulate e.g. the soil environment or root internal states.

We present the use of CRootBox for five different examples. The simplest use of CRootBox, i.e., growth of single root systems in unconfined space in homogeneous soil is demonstrated based on 22 different sets of previously published root architectural parameters (Example 1 in Fig. 1). Root growth in confined containers as a way to mimic experimental setup is demonstrated in Example 2. Example 3 shows a simulation with more than 200 individual root systems to represent a field plot. Virtual soil coring is performed; such data could be compared to results of classical field sampling methods. Chemotrophic root growth is shown in Example 4. To exemplify coupling with an external soil model, we present a virtual case study in Example 5 in which we simulate a growing root system in a soil core under irrigation. The underlying soil model is taken from Bittelli *et al*. (2015) and the related code downloaded from http://www.dista.unibo.it/~bittelli/soilphysicspython.php. We merged our root architecture model into this code in order to demonstrate how such a coupling works. As part of that coupling example, we additionally provide a Python code for the numerical solution of the Doussan model for water flow inside the root system (Doussan *et al*. 1998).

The code of the CRootBox model, as well as its different applications is available at the address: https://plant-root-soil-interactions-modelling.github.io/CRootBox/

Finally, we provide a discussion on the above examples, the potential of the proposed framework and future perspectives. We will demonstrate specifically 1) that CrootBox enables the modelling of mature root systems of a large range of plant species, 2) it enables root system modelling at the field scale, 3) it enables extension to the whole plant system and 4) it facilitates an easy and straightforward communication with environmental models.

## 2 Materials and Methods

### 2.1 CRootBox model

CRootBox is an advanced re-development of the root architecture model RootBox (Leitner *et al*. 2010a). It was translated into C++ and thus the structure was changed from L-Systems to an object oriented design. In the following paragraphs we will give a short summary of the processes that are included in the model, describe model extensions for mature root systems, and then, the basic object oriented layout, and the interface for root function modelling.

#### Topology

The development of the root system is described by the growth of individual roots having specific root types that are governed by model parameters for each type. The production of successive lateral roots may follow root system topology. Alternatively, there could be several possible successor root types, each with a certain probability (Pagès *et al*. 2004). This is defined by the parameter *successor*. Table 1 presents a complete list of parameters including units. For the definition of a root system, the parameters must be determined for each root type.

**Table 1.**
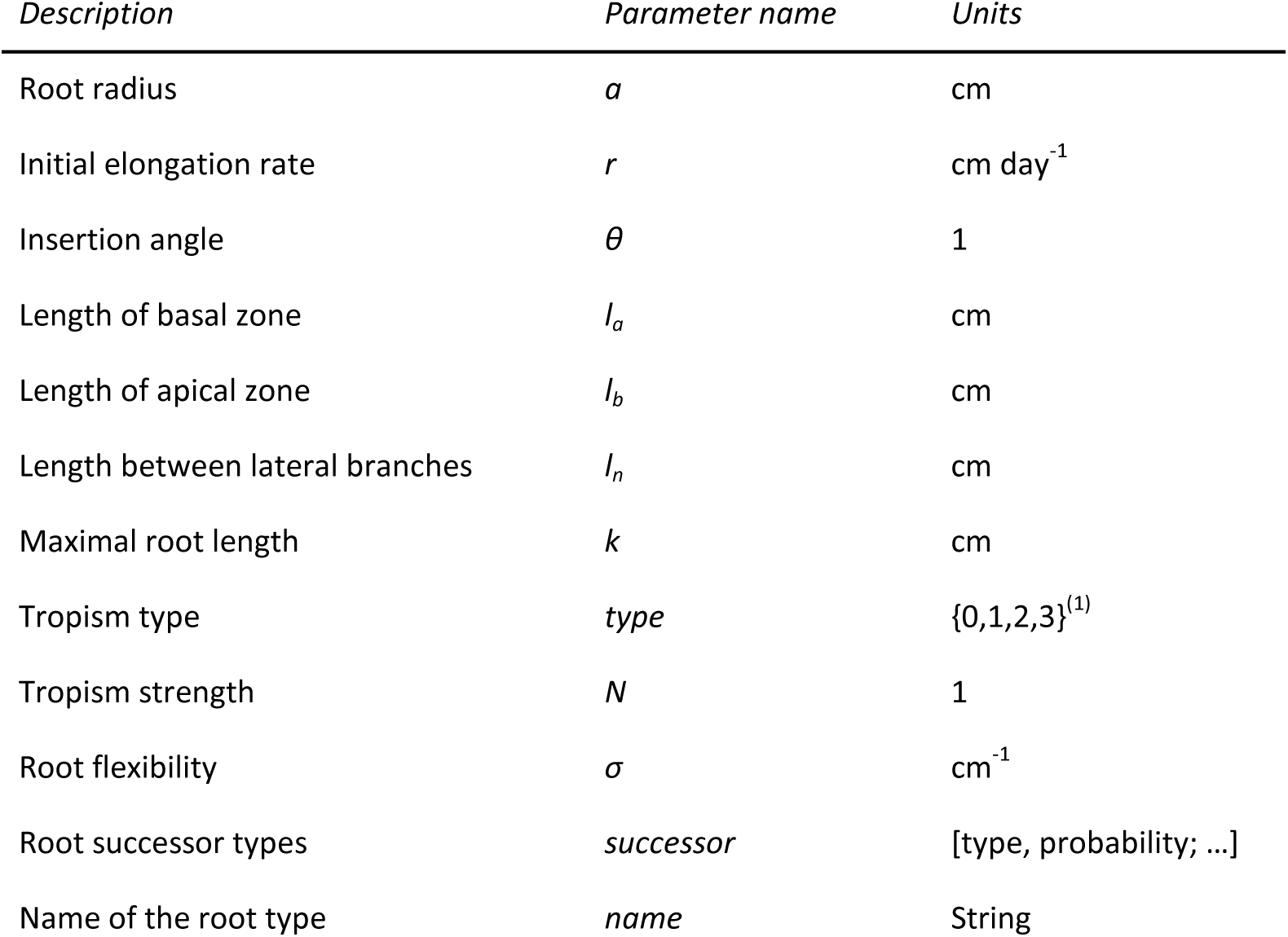
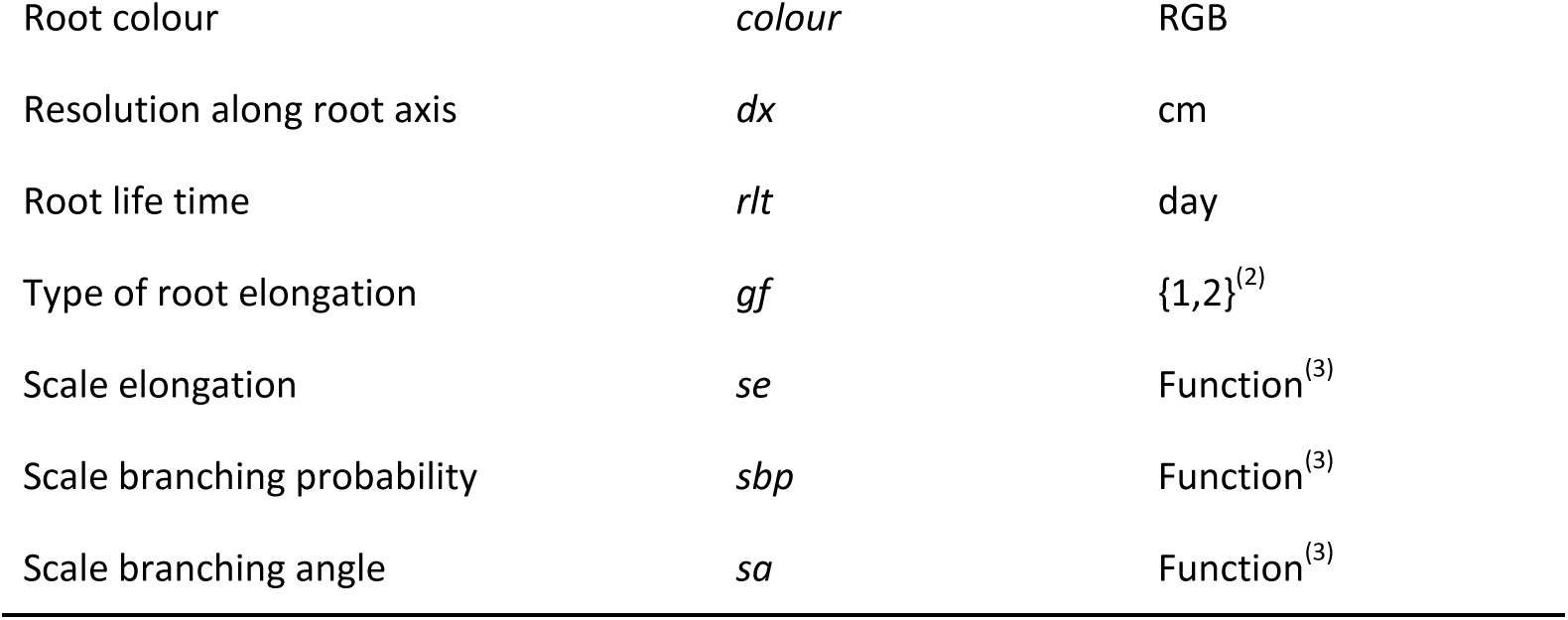
Complete list of parameters used by CRootBox for each root type. ^(1)^ Predefined tropism types are plagio-, gravi, exo-, chemo-or hydrotropism. ^(2)^ Predefined types are negative exponential and linear growth. ^(3)^ Predefined callback functions depending on a soil property to realize root responses

#### Growth

Each individual root elongates as long as the root age is smaller than the root life time *rlt*. The length of the root at a certain time *t* is given by linear growth *l_lin_* or negative exponential growth *l_exp_*,

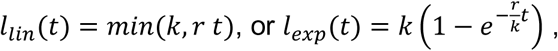

where *k* is the maximal length, and *r* is the initial growth rate.

Each root with laterals is divided into a basal zone, a branching zone, and an apical zone. After the basal zone and the apical zone have developed, lateral roots start to emerge with a fixed branching angle *θ*. The maximal root length *k* of a root is given by

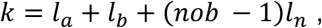

where *l_a_* is the length of the apical zone, *l_b_* is the length of the basal zone, *l_n_* is the interbranching distance and *nob* is the maximal number of laterals the root can develop.

#### Tropism

A change in direction of the growing root tip occurs every distance *dx*, which is the axial resolution of the root. After each distance *dx*, the root tip orientation is randomly changed to represent soil tortuosity. For directed trophic growth, the change in direction of root tip is calculated according to a random optimisation process. We randomly choose *N* rotational changes in growth direction and pick the one that minimises an objective function. This objective function defines the type of tropism that is described, e.g. gravitropism picks directional changes that are downwards, or hydrotropism, a response of root growth to gradients in soil water content. Therefore, the tropism is described by three parameters: *type* defines the objective function, *N* the number of trials, and *σ* is the flexibility of the root, i.e. the strength of change in root direction.

Additional parameters are the name of the root type (*name*), and the root colour (*colour)*. Furthermore, three callback functions can be defined to realise root responses to the soil environment in terms of elongation rate (*sef*), branching density (*sbpf*), and branching angle (*saf*). These functions are described in Section 2.4. All root type parameters except *name, colour, sef, sbpf, saf*, and *successor* are given by mean and standard deviation.

Note, that the overall root architectural growth is computed using a recursive algorithm that can give output at any specified time points, i.e., no forward loop with explicit time steps is required for the root growth modelling. Time steps are only necessary in the framework of a split operator-type coupling with another model such as a soil model.

### 2.2 Modelling mature root systems

In grown root systems, different physiological types of roots can be distinguished. In CRootBox, we follow the nomenclature of the international society for root research, ISRR (Gregory 2006; Zobel and Waisel 2010):

1. Tap root: The first root that emerges from the seed.
2. Lateral roots: First order laterals are any roots that branch from tap root, basal roots, or shoot borne roots. Second order laterals are lateral branches from first order laterals.
3. Basal roots: Emerge from the hypocotyl or mesocotyl. In literature they are also referred to as seminal roots in monocotyledon plants.
4. Shoot borne roots: Emerge from shoot tissue. In literature they are also referred to as adventitious roots, nodal roots, or crown roots.

In dicotyledonous plants root types (1)–(3) are present, see Fig. 2(a). Shoot borne roots are not formed in dicotyledonous plants (Chochois et al. 2012; Hochholdinger et al. 2004). To model dicotyledonous plants in CRootBox, the emergence times of the basal roots must be specified. This is done by three parameters: the first describes the occurrence of the first basal root *first_B_*, the second the time delay between the emergence of basal roots *delay_B_*, and the third the maximal number of basal roots *max_B_*.

**Fig. 2.**
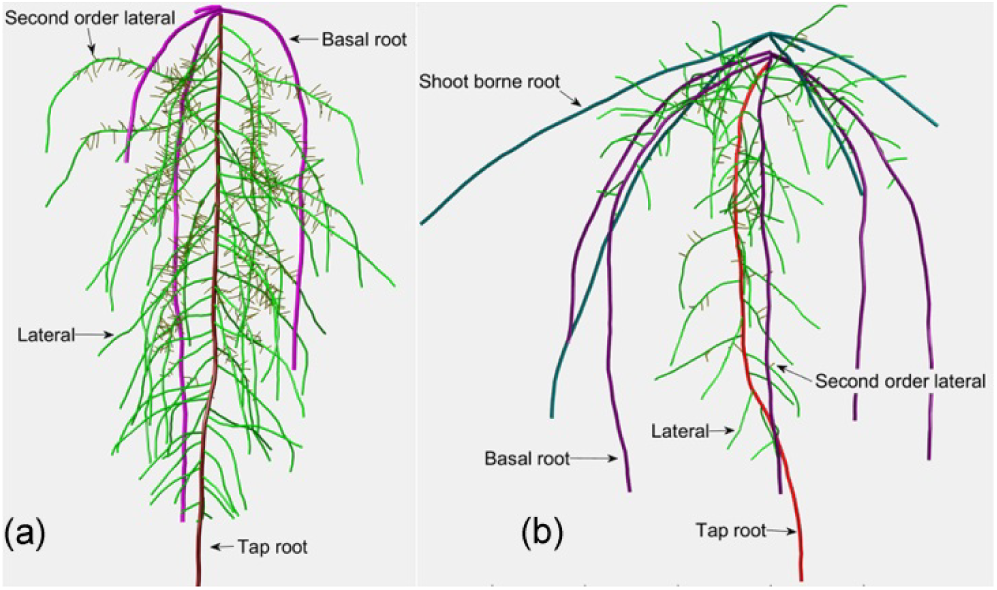
ISRR nomenclature for a dicotyledon plant (a) and a monocotyledon plant (b). Root colours denote the different root types: tap root (red), basal root (magenta), shoot borne root (cyan), lateral root (green), second order lateral (yellow)

In monocotyledonous plants all root types (1)-(4) can be observed, see Fig. 2(b). To model a monocotyledon plant the emergence of basal roots is described by *first_B_*, *delay_B_*, and *max_B_* as in the dicotyledonous case. Additionally, the shoot borne roots are described by four parameters following Klepper (1991): The occurrence time of the first shoot borne root is denoted as *first_S_*. The time delay between successive shoot borne roots is called *delay_S_* and is related to the phyllochron. The number of shoot borne root axes per root crown is named *n_s_*, and the vertical distance between root crowns *dz_S_*. The angle between the root axes along a single root crown is defined as *2π*/*n_s_*.

The planting depth is given by the parameter *depth*. Hypocotyl and mesocotyl are not simulated explicitly. The location of the hypocotyl is assumed to be between the soil surface and the planting depth (*depth*). The location of the mesocotyl lies between half of the planting depth and the seed. Basal roots emerge at the seed, and the first shoot borne root emerges above the mesocotyl. Successive root crowns move vertically up the plant shoot. Table 2 summarizes the plant parameters and their units.

**Table 2.** List of plant parameters needed for the root architecture development of dicotyledonous and monocotyledonous plants

| *Description* | *Parameter name* | *Units* | *Plant type* |
| --- | --- | --- | --- |
| Planting depth | *depth* | cm | Dicot and monocot |
| First emergence of basal roots | *first<sub>B</sub>* | day | Dicot and monocot |
| Time period between basal roots | *delay<sub>B</sub>* | day | Dicot and monocot |
| Maximal number of basal roots | *max<sub>B</sub>* | 1 | Dicot and monocot |
| First occurrence of shoot borne roots | *first<sub>S</sub>* | day | Monocot |
| Time period between shoot borne roots | *delay<sub>S</sub>* | day | Monocot |
| Number of shoot borne roots per root crown | *n<sub>S</sub>* | 1 | Monocot |
| Distance between root crowns along the shoot | *dz<sub>S</sub>* | cm | Monocot |

### 2.3 C++ structure of CRootBox

The object oriented model structure uses the principle of code reuse and encapsulation, in order to make the code easier to understand and use. Therefore, the root architecture is described by meaningful objects that interact with each other. The structure of the CRootBox framework is outlined in Fig. 3. An in-depth description is given in the doxygen class documentation (see supplementary data S1).

**Fig. 3.**
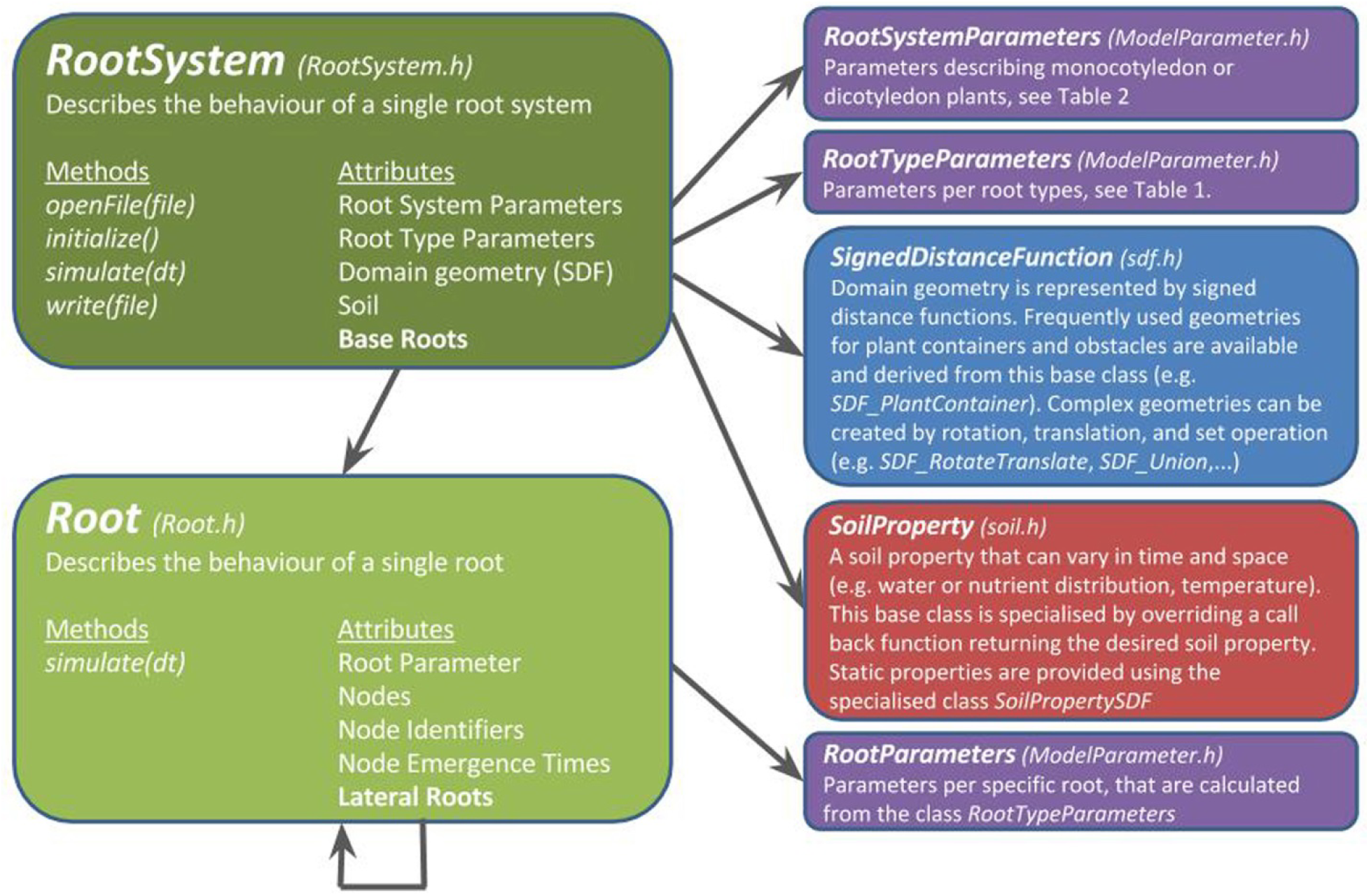
The most important classes of CRootBox including a short description, and the main methods and attributes of the classes *Root* and *RootSystem*. Colours represent different header files.

The simulation itself is performed by the class *RootSystem*, which describes a single root system and manages (1) all root system parameters, (2) the base roots of the system, i.e. the tap root, basal roots, and shoot borne roots, (3) domain geometry, i.e. confining geometries and obstacles, and (4) offers utility functions for basic analysis of results, and extensive output functionality for visualisation and analysis.

Model parameters are represented by three classes: The two classes *RootTypeParameters* and *RootSystemParameters* exactly mimic the parameters given in Table 1 and Table 2. Additionally, the class *RootParameters* stores the parameters for a specific root, i.e. a single realisation of the values from *RootTypeParameters* that are given by mean and standard deviation. The development of the base roots is determined by these parameters and described by the class *Root*, which recursively manages all its lateral roots which are also of class type *Root*.

Simulation results can be exported and analysed by the class *RootSystem*. For visualization with ParaView, root architecture can be represented in the VTK Polygonal Data format (VTP), and the domain geometry in a ParaView Python script. Simulation results can be exported to the root system markup language RSML (introduced by Lobet *et al*. 2015), which is perfectly suited for comparison with experimental measurements. Furthermore, DuMu^X^ DGF format can be used to calculate porous media flow problems within and around the root geometry. And finally, results can be exported as plain text for general analysis (e.g. Python, R, or Excel). Analysis includes auxiliary functions to retrieve resulting states from the individual roots (e.g. age, length, radius). Furthermore, the topology of the root system can be easily retrieved by the methods *RootSystem::getNodes* and *RootSystem::getSegments*. These two methods can be used to build an adjacency matrix of a mathematical graph that represents the root system. This strongly promotes the implementation of root internal models. In Section 3.5 we demonstrate this approach by presenting the calculation of the xylem flux following Doussan et al., 2006. Model equations and numerical derivation are presented in Appendix A.

Post-processing in C++ is facilitated by the class *SegmentAnalyser*. While *RootSystem* offers analysis tools per root, the class *SegmentAnalyser* works per segment. Main features include depth distributions of arbitrary parameters (e.g. root length, or root surface distributions), cropping with a geometry given by a signed distance function (see Fig. 11 (b) for an example), and thresholding of arbitrary parameters. These tools were developed to mimic general experimental procedures, like for example soil coring, or the analysis of soil trenches. While C++ is generally not well suited for post-processing, it is an advantage to have these steps predefined directly in C++, as it is an enormous speed up compared to other software like Python, R, or Excel.

The code and Doxygen documentation are available in the github reposirory https://plant-root-soil-interactions-modelling.github.io/CRootBox/.

### 2.4 Describing root responses to environmental conditions

CRootBox is the only root architecture model which is independent of time step and axial resolution. This means the resulting root length, number of segments and horizontal and vertical spread of the root system will be the same, if the overall simulation time is partitioned into months or into seconds. Furthermore, overall root length will not differentiate choosing small or large segment lengths as axial resolution. This makes CRootBox very suitable for creating root functional models that can describe processes at different temporal and spatial scales.

Root responses to soil properties are implemented with a generic approach: A scalar soil property is described by the class *SoilProperty* that provides a lookup method (*SoilProperty::getValue*) for the parameter value which can vary in space and time. Soil properties can be any scalar value, for example water content, nutrient concentration, soil strength, microbial activity, or temperature. The class *SoilProperty* is used to

1. describe hydro- or chemotropism, which is implemented in the same way as in Leitner et al. (2010a). Briefly, it is based on a random optimisation towards the largest gradients of water content or solute concentration between the current and projected next position of the root tip.
2. alter the root elongation rate. This is realized by a new parameter ‘scale elongation’ (*se*), see Table 1. Therefore, the root elongation rate is scaled by the value returned by *SoilProperty::getValue*. A value smaller than 1 leads to impeded growth, a larger value to enhanced growth.
3. scale the root branching angle θ. This is realized by a new parameter ‘scale angle’ (sa). A value smaller than 1 leads to more acute angle, a larger value to a more obtuse angle.
4. scale the root branching density. Branches potentially emerge at a given internodal distance *ln*, which is the minimal possible distance between the laterals. The function ‘scale branching probability’ (*sbp*) can be used to reduce the branching density.

### 2.5 Root growth inside containers

To mimic experimental settings it is important to precisely represent plant containers and obstacles. The domain geometry is realised using signed distance functions (Osher and Fedkiw 2003) that are represented by the class *SignedDistanceFunction*. These functions return the distance to the closest boundary for each point in the domain. A negative value refers to a point that is inside a given domain, a positive value to a point that is outside. *SignedDistanceFunction* is the base class of all such geometries (e.g. *SDF_PlantContainer)*. More complex geometries can be easily created by rotating and translating a base geometry using the class *SDF_RotateTranslate*, and by using set operations like union (*SDF_Union*), difference (*SDF_Difference*), or intersection (*SDF_Intersection)*.

If the new root tip position of a growing root does not lie within the geometric boundaries, a new pair of insertion and radial angle, (α, β), is chosen as follows: First, only β is chosen uniformly random between -π and π while α is left unchanged. If, after a maximal number of trials, no new valid pair α and β has been found, α is increased by a small increment, and the procedure for finding an angle β starts again. This simple approach leads to realistic root behaviour at the boundaries, where thigmotropism can be observed.

### 2.6 Add-ons to CRootBox

Database of root architectural model parameters

Based on literature sources that published root architectural parameters and that are suitable for modelling, we created a database of 22 parameter sets for 14 different species. If the parameters were published for another model and needed to be adapted to the requirements of CRootBox, we computed them, if possible, based on available data and otherwise estimated the value such that the resulting root system was as similar as possible to the original one by visual comparison. Details are presented in Appendix B.

The parameter sets are available at the address: https://doi.org/10.6084/m9.figshare.c.3745478.

#### Web application

Visualisation of the CRootBox capabilities for the growth of single root systems is available through a web application at https://plantmodelling.shinyapps.io/shinyRootBox. The user chooses a parameter set, gets information about the underlying publication that contains the model parameters, and, upon pressing “Unleash CRootBox”, receives a 3D visualisation of the newly generated root system as well as standard metrics such as root length density profiles, number of roots and number of segments. Parameter values can be changed interactively and the results immediately visualised. The 3D root systems can be stored in VTP, CSV and RSML formats and thus offers a quick and easy opportunity for the creation of single root systems.

#### Python binding

The CRootBox code is fast and easy to read for everyone who is used to work with C++. However, we felt that the use of CRootBox should not be limited to this group of persons. Therefore, we created a Python binding using the C++ library Boost.Python. After the CRootBox code has been compiled once using this class, all the exposed classes and methods can be used in simple Python scripts that are very similar to the previous Matlab scripts of Rootbox (Leitner *et al*. 2010a), however still perform with the speed of a C++ code.

Furthermore, Python has evolved to be a universal gluing language for coupling different pieces of software (e.g. OpenAlea, Pradal *et al.* 2008). With the Python binding of CRootBox it is directly possible to create root systems and use those with any Python-based soil code. We demonstrate this in Section 3.5 by using the code “PSP_infiltrationRedistribution1D” from the book “Soil Physics with Python” (Bittelli *et al*. 2015), adding a sink term for root water uptake based on CRootBox simulated root architectures. We want to emphasize that Python makes it very easy to couple to any soil models which often are solved by partial differential equation (PDE) solvers like Comsol, DuMu^x^, or specialized solutions that are available in C++, Matlab, or Python.

Fig. 4 summarises the available features of CRootBox. The core C++ code can be used as a stand-alone model and offers the highest level of flexibility. The Python library exposes the main functions and variables of CRootBox so they can be used in much easier Python scripts. Python is also increasingly important as gluing language for model coupling (Perez *et al*. 2011). The web application features the basic capabilities of CRootBox in a graphical user interface online and offers access to the data base of root architectural models which we compiled based on literature sources. Thus it offers an opportunity to quickly create virtual root systems of single plants and store them in different data formats.

**Fig. 4.**
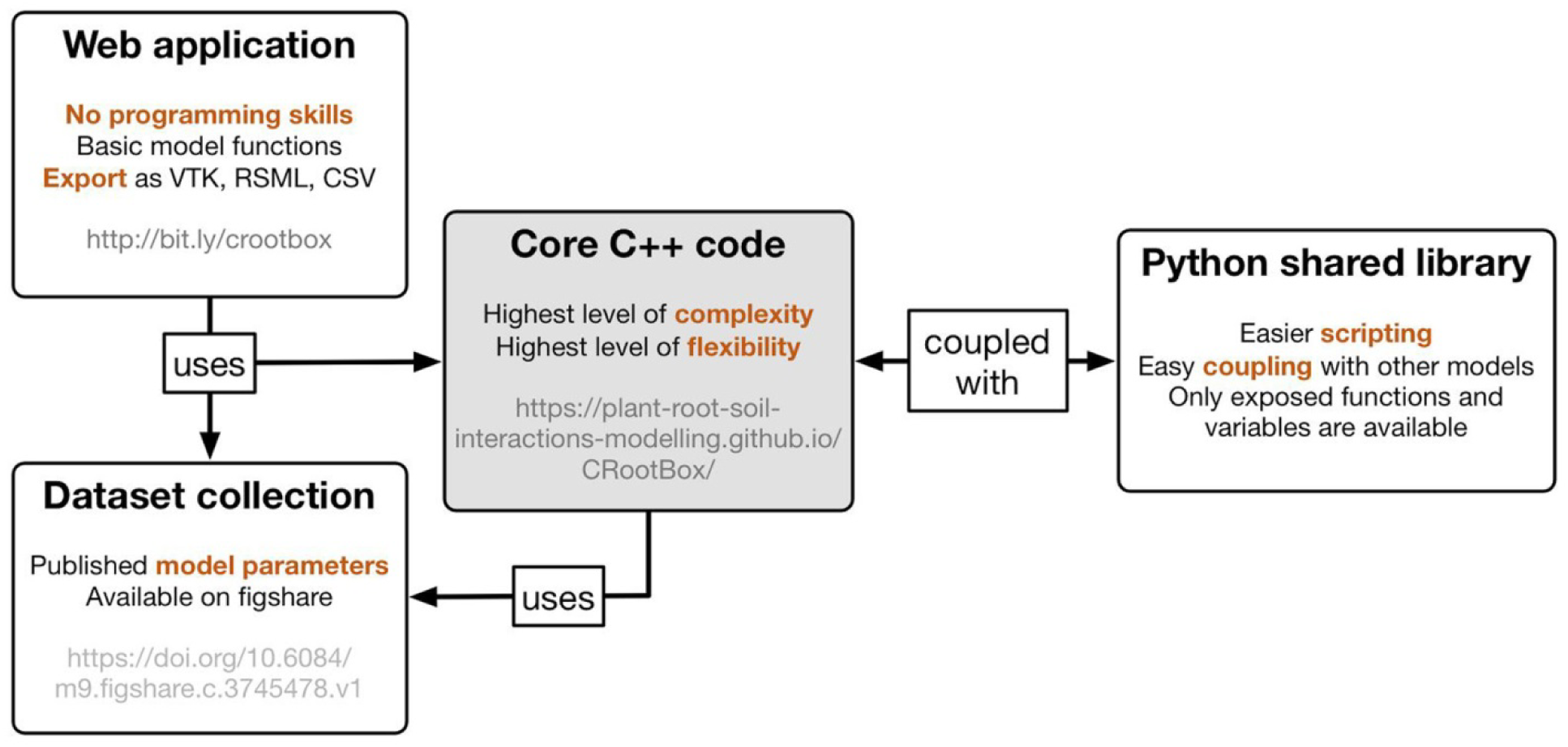
CRootBox is presented through its core C++ code as well as three add-ons that simplify its use for specific purposes: the dataset of root architectural parameters, the web application for simulation of single root systems and export of related structures, and the Python shared library for simpler scripting and coupling to external models.

## 3. Results

### 3.1 Example 1: Unconfined growth of individual root systems

In Fig. 5, we show visualisations of the 22 different root system parameter sets currently stored in our database, after a simulation period of 8 weeks. Fig. 5 shows that CRootBox is capable of simulating a wide variety of different types of root systems, including fibrous and tap root systems.

**Fig. 5:**
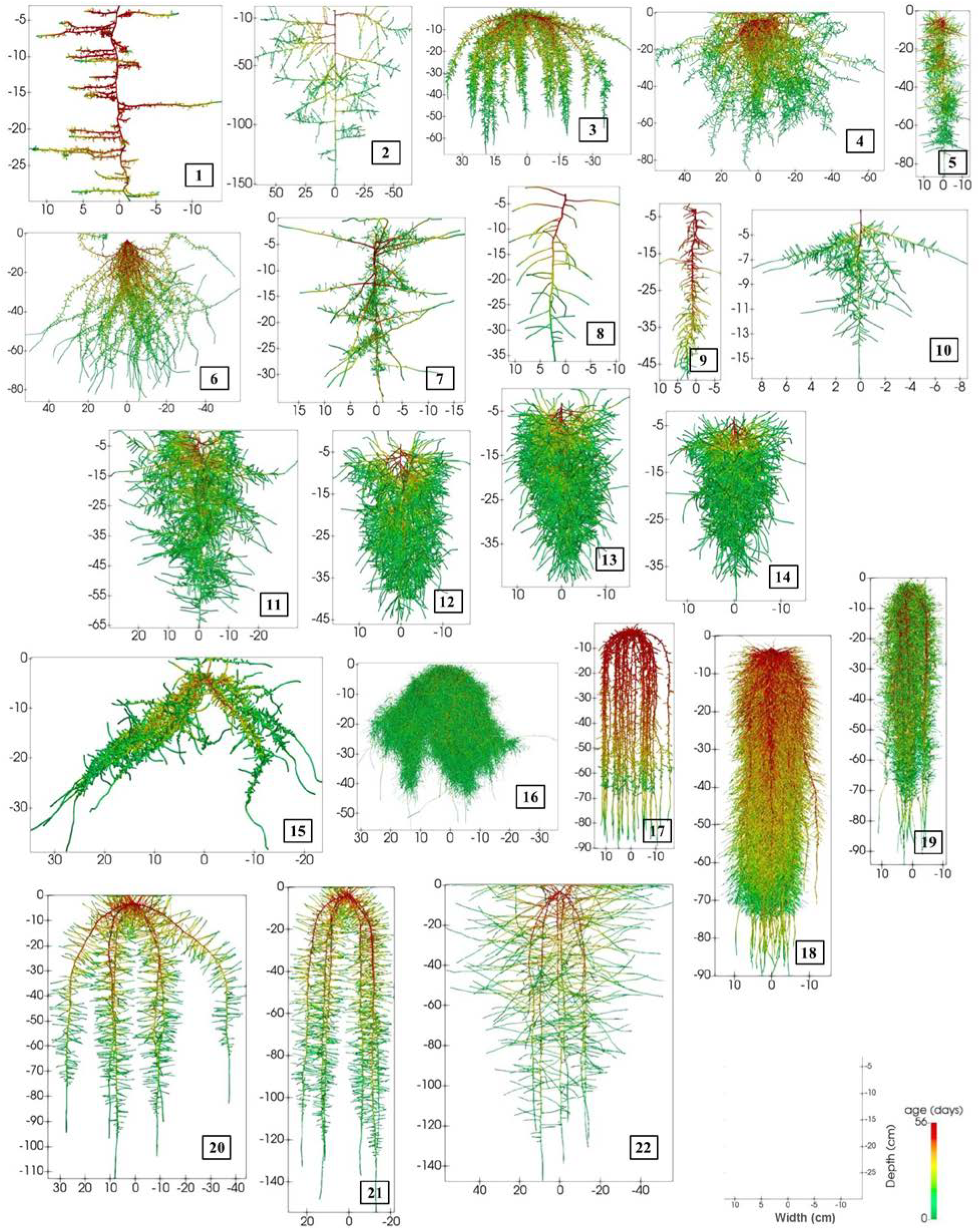
3D visualisation of the root systems simulated with unconfined growth based on the parameters contained in the database (1. *Anagallis femina* 2. *Brassica napus* 3. *Brassica oleracea* 4. *Crypsis aculeata* 5. *Helianthus* L. 6. *Juncus squarrosus* 7. *Lupinus albus* 8. *Lupinus angustifolius* 9. *Medicago truncatula* 10. *Noccaea caerulescens* 11. *Pisum sativum* a 12. *Pisum sativum* b 13. *Pisum sativum* c 14. *Pisum sativum* d 15. *Sorghum bicolor* 16. *Triticum aestivum* 17. *Zea mays* 1 18. *Zea mays* 2 19. *Zea mays* 3 20. *Zea mays* 4 21. *Zea mays* 5 22. *Zea mays* 6). For visualization purposes, the root radii on average were increased five-fold.

Simulation outcome is the full 3D geometry of the root system. In some cases, more aggregated information is required for further analysis or for use in simpler models that could not handle 3D root architectural information. Furthermore, CRootBox is a stochastic model in which each parameter is defined by its mean and standard deviation. Thus, each simulated root system is only one of many possible realisations of this parameter set. Based on 100 realisations of each of the parameter sets in the database, Fig. 6 shows the mean plus/minus standard deviation of root length distributions (RLD) with depth, by summing up all the lengths of the root segments in 5 cm depth intervals, divided by the layer thickness, thus giving units of cm root length per cm of soil. The resulting root length distributions vary strongly between the different datasets, maximal value of the RLD of fibrous root systems ranging between 400 and 1000 cm cm^-1^, those of tap root systems ranging between 20 and 80 cm cm^-1^. The standard deviation of the RLD depends on the standard deviations of the different model parameters and may thus vary considerably: For published root architectural parameters, this information is not always provided, in which case we set the standard deviation to 10% of the mean value. The dynamic development of selected root systems and its corresponding RLD profiles are presented in Figs. 7 and 8 for a tap and fibrous root system, respectively. Field scale simulations and subsequent virtual coring allows comparison with coring data available in literature. This will be described in section 3.3. The following section describes simulations that mimic different experimental setups where plants are grown in confining containers.

**Fig. 6:**
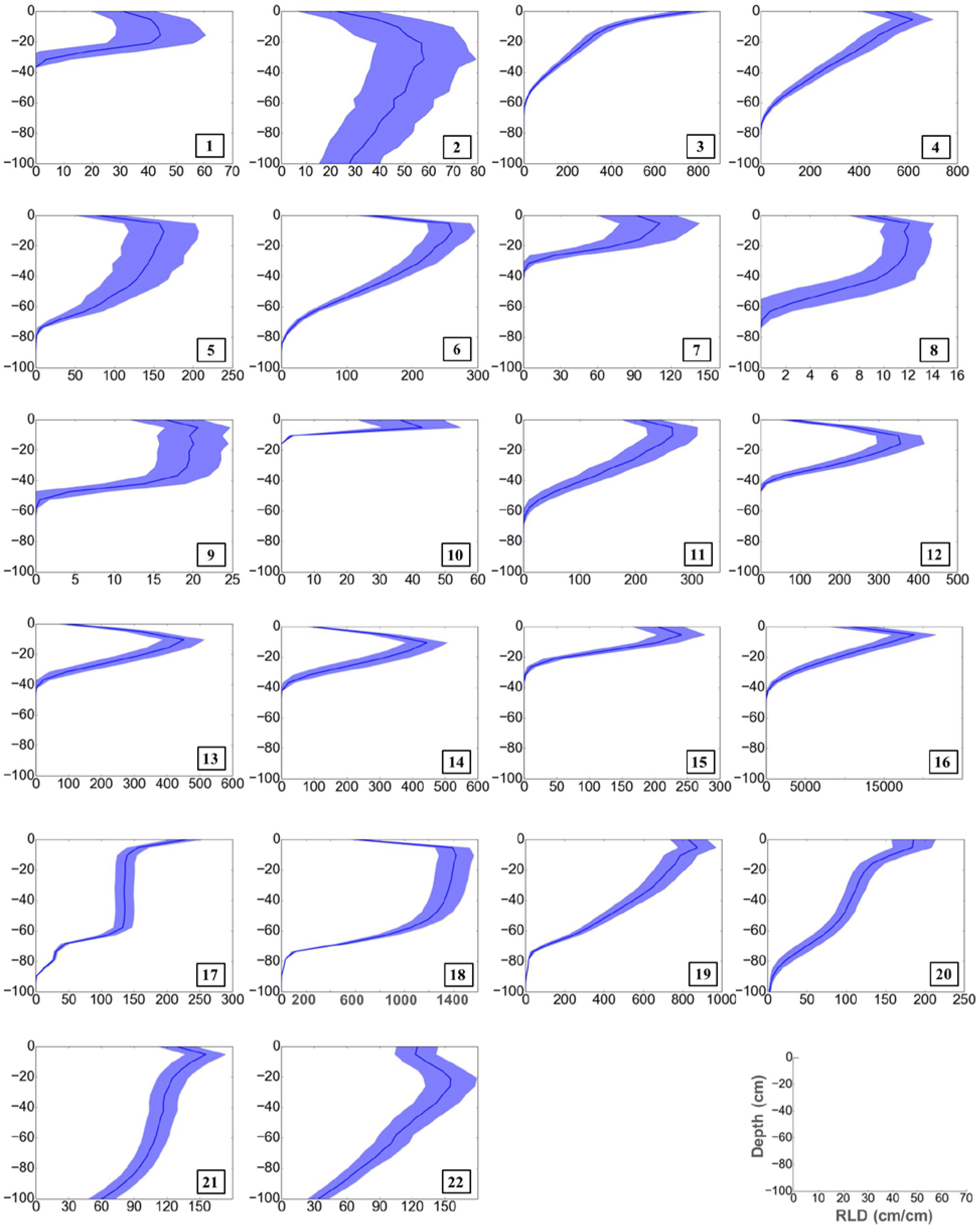
Root length distributions with depth (cm root length per cm soil depth) corresponding to the root systems of Fig. 1; represented by the mean (dark blue line) and plus/minus standard deviation (light blue bands) of 100 realisations (1. *Anagallis femina* 2. *Brassica napus* 3. *Brassica oleracea* 4. *Crypsis aculeata* 5. *Helianthus* L. 6. *Juncus squarrosus* 7. *Lupinus albus* 8. *Lupinus angustifolius* 9. *Medicago truncatula* 10. *Noccaea caerulescens* 11. *Pisum sativum* a 12. *Pisum sativum* b 13. *Pisum sativum* c 14. *Pisum sativum* d 15. *Sorghum bicolor* 16. *Triticum aestivum* 17. *Zea mays* 1 18. *Zea mays* 2 19. *Zea mays* 3 20. *Zea mays* 4 21. *Zea mays* 5 22. *Zea mays* 6). Due to the large variation of RLD, especially between fibrous and tap root systems, the x-axis scale differs for the different root systems.

**Fig. 7:**
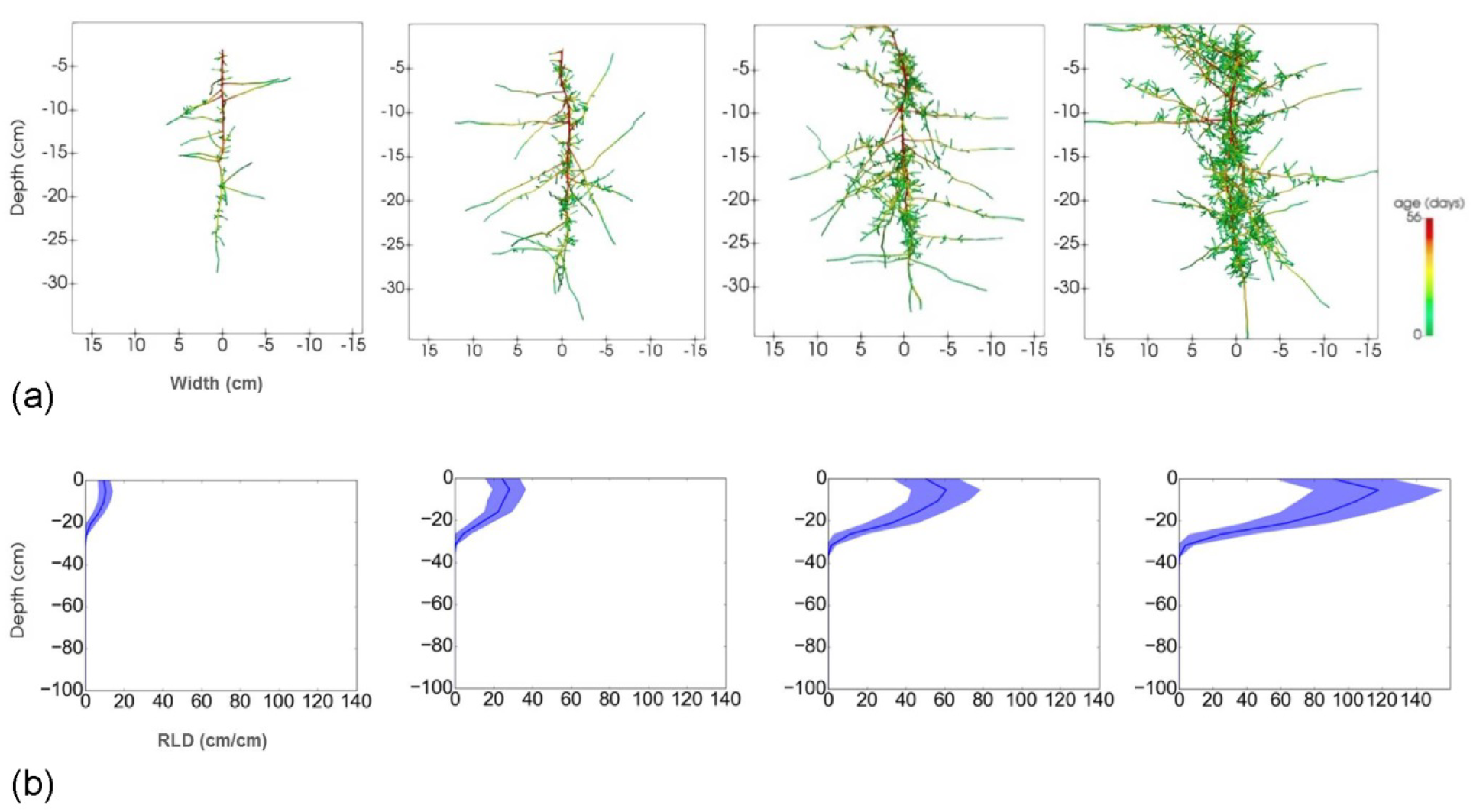
*Lupinus albus* root system after 14, 28, 42 and 56 days. (a) 3D visualisation and (b) corresponding root length distributions with depth (cm root length per cm soil depth) represented by mean (dark blue line) and plus/minus standard deviation (light blue bands) of 100 realisations.

**Fig. 8:**
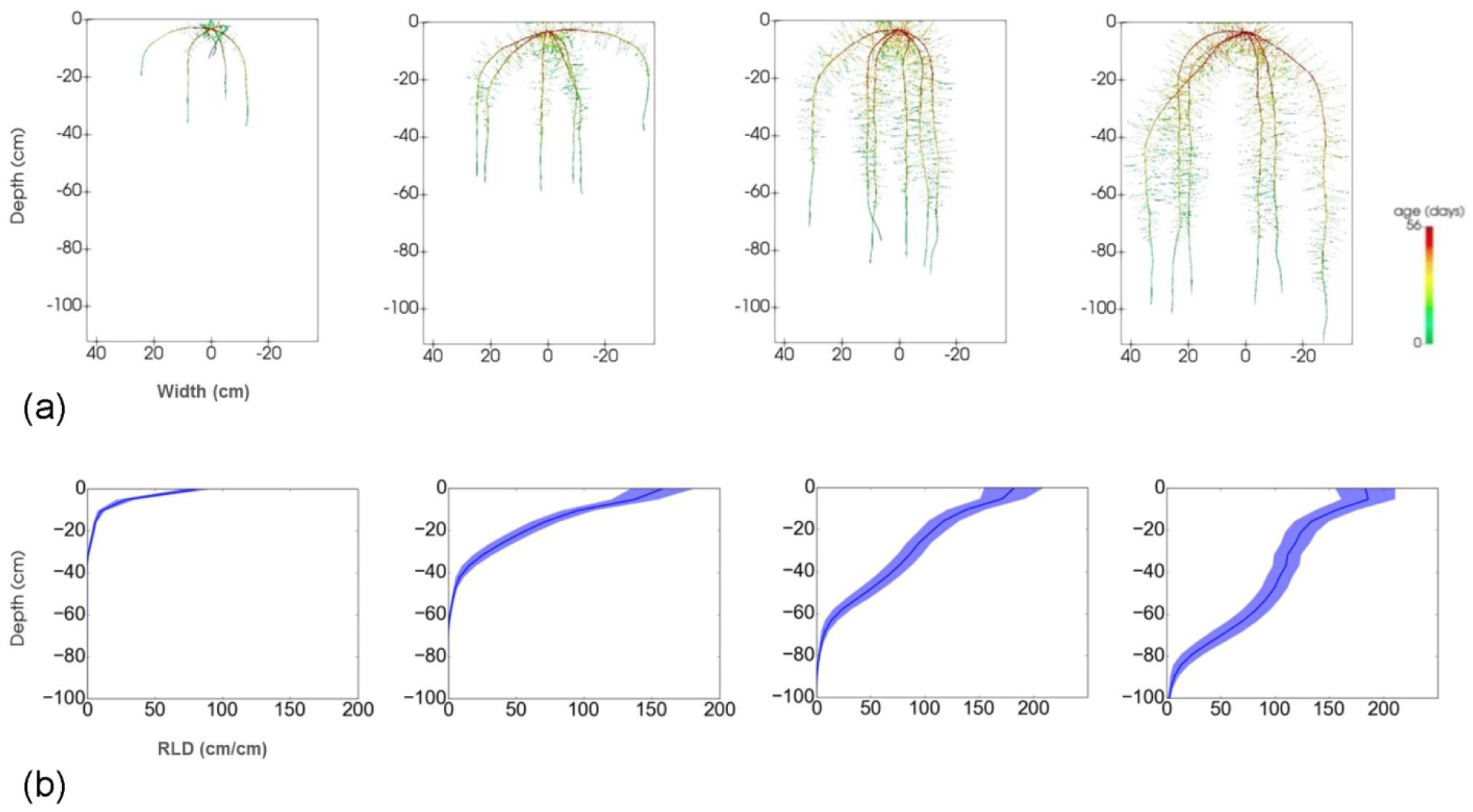
Root length density of *Zea mays* root system after 14, 28, 42 and 56 days. (a) 3D visualisation and (b) corresponding root length distributions with depth (cm root length per cm soil depth) represented by mean (dark blue line) and plus/minus standard deviation (light blue bands) of 100 realisations.

### 3.2 Example 2: Confined growth

Root systems can be grown virtually in containers using CRootBox, e.g., in order to mimic experimental setups like pot or rhizotron experiments. Predefined containers include pots of cylindrical or conical shape as well as rhizotrons, i.e., rectangular containers, which can be set at a user-defined inclination. However, virtually any shape can be created using the build-in signed-distance function operators. The root systems of a *L. albus* and a *Z. mays* plant, respectively, growing in a cylindrical pot and in a rhizotron are demonstrated in Figs. 9 and 10, together with the corresponding root length distributions with depth. The simulation time was 56 days, and during this time the roots reached the side and bottom of the container. This is also reflected in the RLD profiles.

**Fig. 9:**
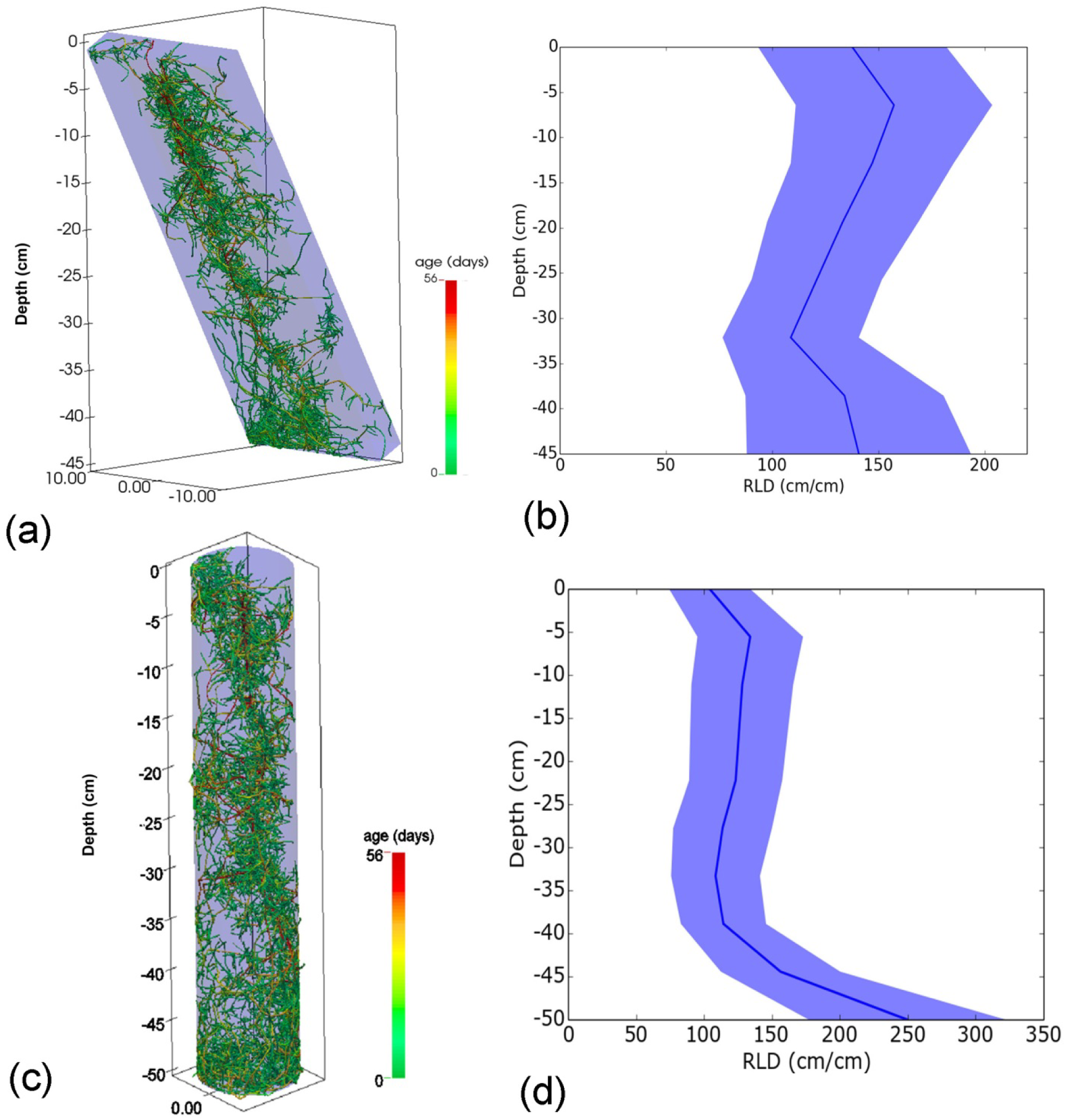
Sample simulation for confined growth of *L. albus*. (a) 3D visualisation of root growth in a rhizotron. (b) Corresponding root length distributions with depth (cm root length per cm soil depth) represented by mean (dark blue line) and plus/minus standard deviation (light blue bands) of 100 realisations. (c) 3D visualisation of root growth in a pot (d) Corresponding root length distributions with depth (cm root length per cm soil depth) represented by mean (dark blue line) and plus/minus standard deviation (light blue bands) of 100 realisations.

**Fig. 10:**
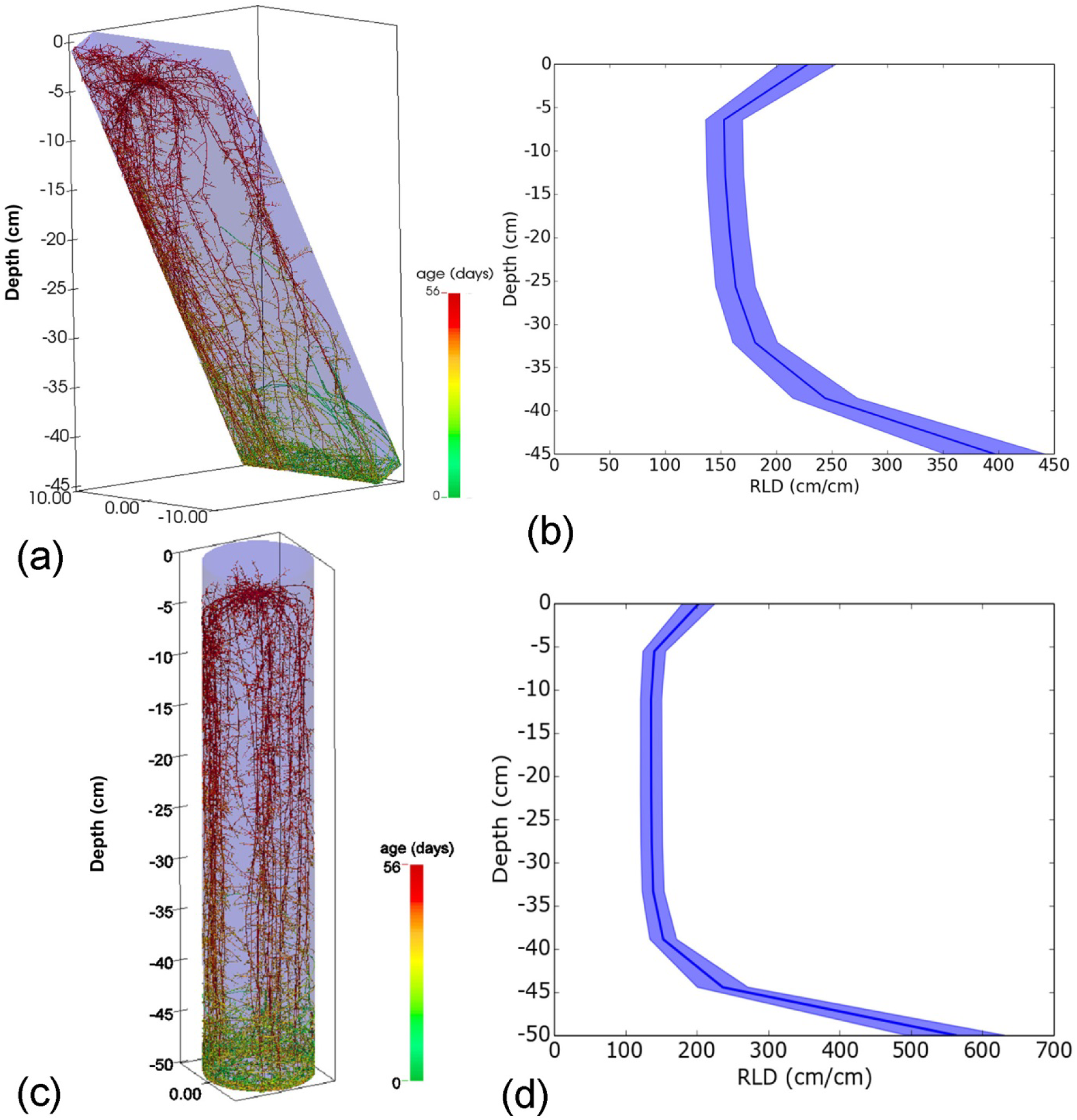
Sample simulation for confined growth of *Z. mays*. (a) 3D visualisation of root growth in a pot. (b) Corresponding root length distributions with depth (cm root length per cm soil depth) represented by mean (dark blue line) and plus/minus standard deviation (light blue bands) of 100 realisations. (c) 3D visualisation of root growth in a rhizotron (d) Corresponding root length distributions with depth (cm root length per cm soil depth) represented by mean (dark blue line) and plus/minus standard deviation (light blue bands) of 100 realisations.

### 3.3 Example 3: Field scale modelling

We simultaneously simulated the 3D root architectures of 222 individual *Triticum aestivum* plants over a vegetation period of 240 days. All the plants were simulated with fifteen main (seminal and crown) roots (Barraclough *et al*. 1989). The model domain size comprised 6 rows each consisting of 37 plants. We used an inter row distance of 18 cm and a fixed distance of 3 cm between two adjacent plants in a row. The planting depth (seed position) was chosen at 3 cm below the soil surface.

Cylindrical cores of 4.2 cm in diameter and 5 cm deep, up to 160 depths were sampled to determine the RLD (cm cm^-3^) of each sampling volume using the feature of CRootBox that allows to crop the root system in any geometry that is given by a signed distance function. Five rows were sampled with three cores virtually taken in-between two plants in each row, and mean and standard deviation of the RLD were computed. The resulting root architectures are visualised in Fig. 11(a). Fig. 11(b) visualises the soil cores together with those roots that lie inside those cores, and Fig. 11(c) shows the corresponding root length density profiles (cm root length per cm^3^ soil) after 30, 60, 90, 120, 150, 180, 210 and 240 days.

**Fig. 11:**
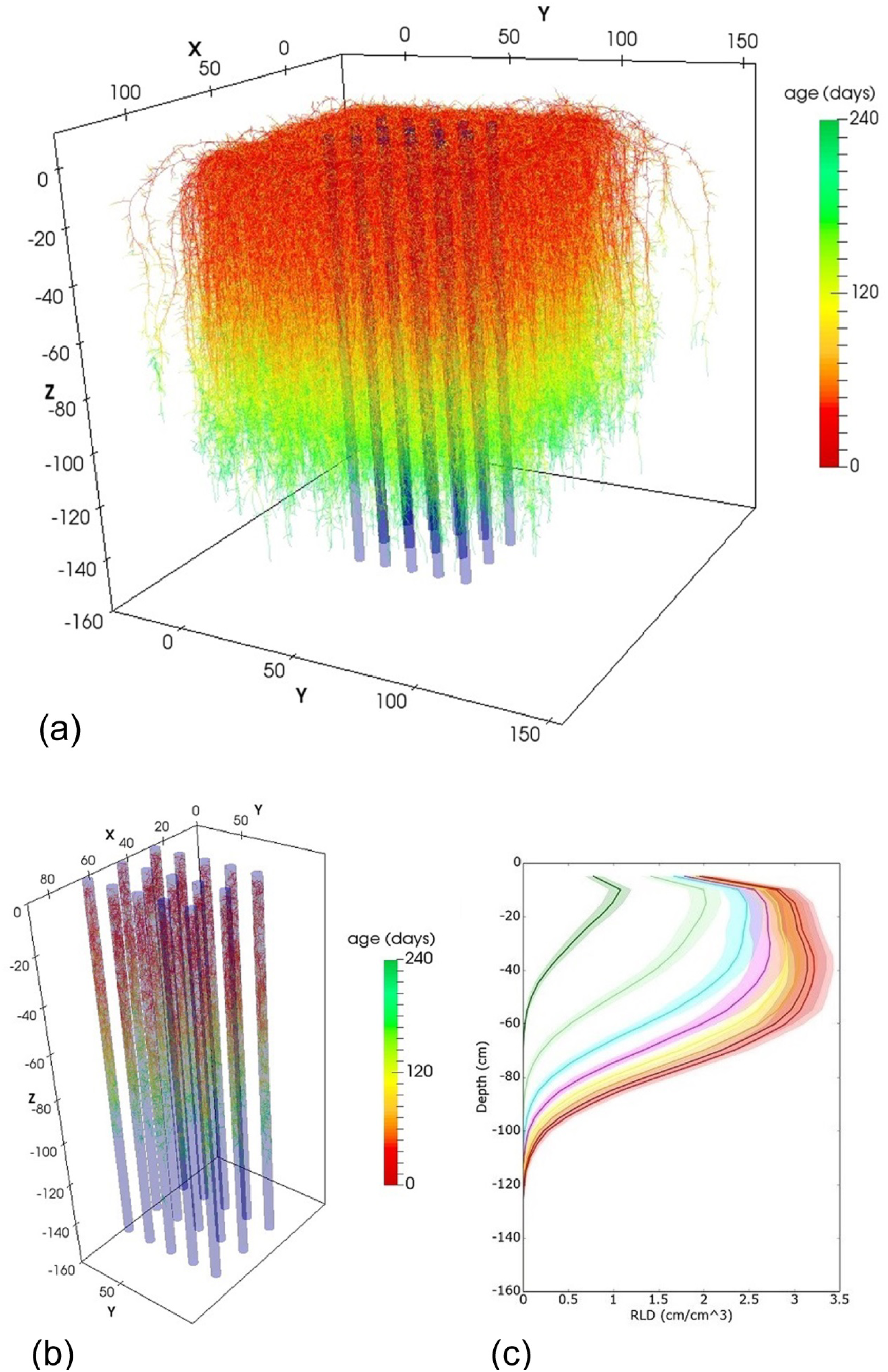
Field scale simulation of *T. aestivum*. (a) 3D visualisation of root growth in a field (b) 3D visualisation of field sampling via coring (c) Root length density profiles (cm cm^-3^) after 30, 60, 90,120,150,180,210 and 240 days, mean and standard deviation from 15 cores.

### 3.4 Example 4: Tropisms

Fig. 12 presents an example of chemotropism, i.e. root growth direction is turned towards locations with higher concentration. In this example, root growth of *Zea mays* follows gravitropism everywhere, and, inside the soil layer or patch with increased concentration, also chemotropism. Both types of tropisms inside the layer or patch are weighted, determining an overall target growth direction. The parameter *N*, which determines the strength of tropism, was set to a value of 3. This value was set the same for each root type in this simulation, but may be specified differently for the different root types if needed. The 3D visualisations in Fig. 12(a,c) clearly show that roots are attracted to stay inside the moist layer or patch, respectively. Fig. 12(b,d) reveal the extent of increased root growth in the soil layer or patch with increased concentration. Such simulations need to be corroborated with experimental data.

**Fig. 12:**
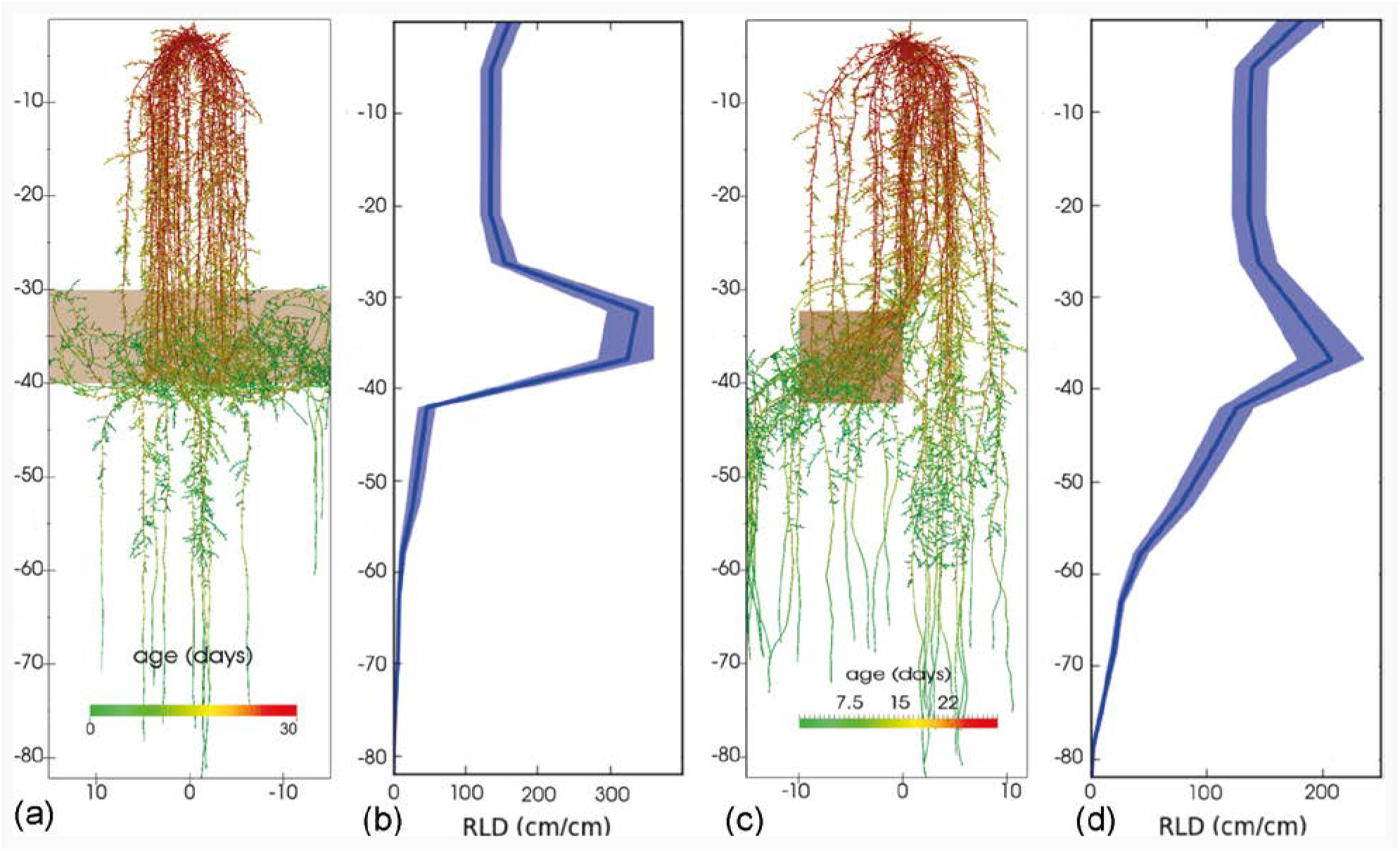
Root growth of *Z. mays* as affected by chemotropism in a soil with a layer or a patch of increased nutrient concentration (*N*=3, *σ*=0.25). (a) 3D visualisation of root growth with chemotropism in a soil with nutrient layer. (b) Corresponding root length distributions with depth (cm root length per cm soil depth) represented by mean (dark blue line) and plus/minus standard deviation (light blue bands) of 100 realisations. (c) 3D visualisation of root growth with chemotropism in a soil with a nutrient patch. (d) Corresponding average root length distributions with depth (cm root length per cm soil depth) represented by mean (dark blue line) and plus/minus standard deviation (light blue bands) of 100 realisations.

### 3.5 Example 5: Coupling to a dynamic soil model, the example from Soil Physics with Python

This is an example to illustrate the coupling of CRootBox with a model of soil water movement (Bittelli *et al*. 2015) and a model of water flow inside the root architecture (Doussan 1998). The soil model of our simulations is based on the code for the solution of the Richards equation from the book “Soil Physics with Python”. For simplicity, we chose the 1D infiltration example *PSP_infiltrationRedistribution1D* (http://www.dista.unibo.it/~bittelli/soil_physics_python.php) for this simulation, however, this can be exchanged with other, more complex models. Root growth was simulated with CRootBox, thereby creating a 3D root architecture. We implemented in Python a numerical solution of the Doussan model; it is given in appendix A. Python was then used as glueing language for each of the three coupled submodels: CRootBox, Richards equation and Doussan model. The root growth and soil and root water flow were solved sequentially at small enough time steps; and water was exchanged, between roots and soil via a sink and source term that depends on the local water pressure gradient inside and outside each root segment. Since the soil model is a 1D model, the sink term for root water uptake from soil was created by averaging root water uptake over all the root segments in each horizontal layer. The coupling is illustrated by the following pseudocode, where *N* is the number time steps, *p*s is the soil water pressure, *p*r is the xylem water pressure:

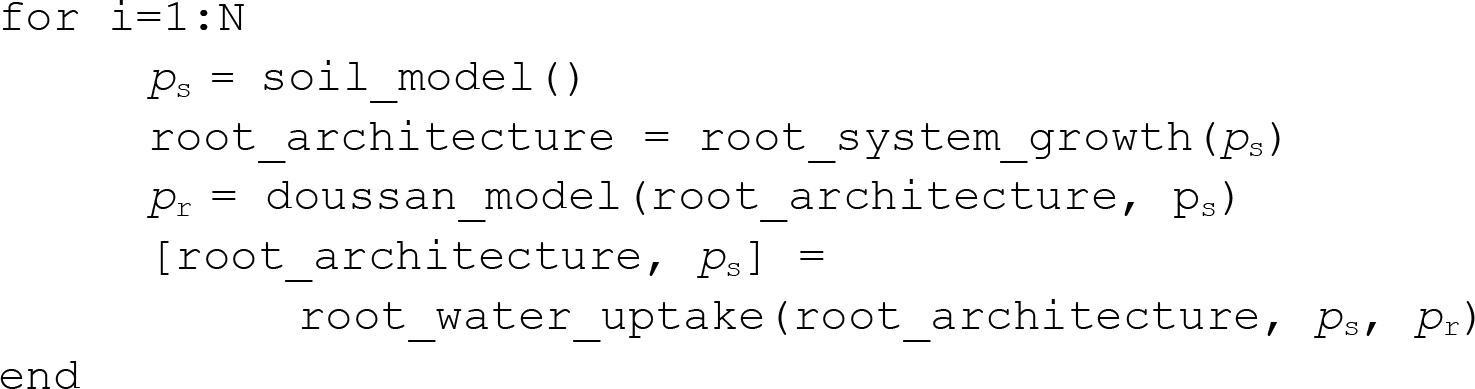

The soil parameters for this simulation were those of a silt loam as provided by the file ‘soilUniform.txt’ that comes together with the Python code of Bittelli et al. (2015); the root architecture was computed with the parameter set for S. *bicolor* from our new database, and the root hydraulic properties were taken from Javaux et al. (2008). The first row shows the root system at day 7, 14 and 21. Colours denote the xylem pressure within the roots. The mid row represents the development of the effective water saturation. Dark areas show the water depletion due to root water uptake. The bottom row shows the root length density (green) and the calculated sink term due to root water uptake (blue), at day 7, 14, and 21. The upper boundary condition for water flow in this example is a dirichlet boundary condition such that there is a constant supply of water from the soil surface. In this example, the soil is moist such that the actual transpiration rate is always equal to the (constant) potential transpiration rate, with *T*_pot_ = -1.15741e-10 m^3^ s^-1^. Thus, there is no water stress, and the integral under the blue curve of Fig. 13 is the same in each time step. We can observe how the sink term follows the root development in this case.

**Fig. 13:**
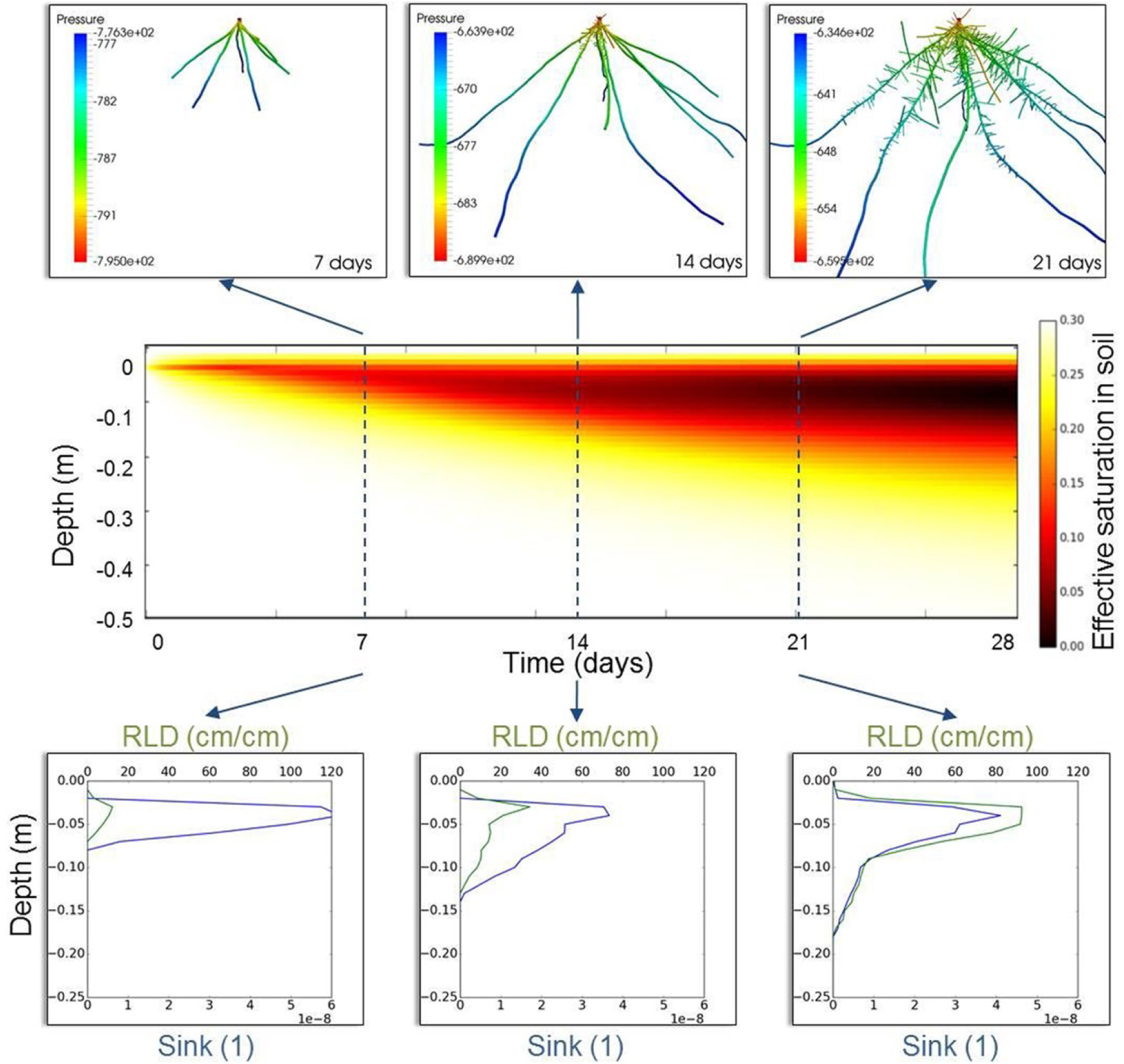
Coupling CRootBox with a 1D Richards Equation solution of “Soil Physics with Python”: The first row shows the root system at day 7, 14 and 21. Colours denote the xylem pressure within the roots. The mid row represents the development of the effective water saturation. Dark areas show the water depletion due to root water uptake. The bottom row shows the root length density (green) and the calculated sink term due to root water uptake (blue), at day 7, 14, and 21.

## 4. Discussion

CRootBox has been advanced from the RootBox model. Improvements include:

1. CRootBox is much faster: While RootBox was restricted to young root systems, CRootBox can easily simulate whole cropping cycles, or even field scale simulations with hundreds of root systems.
2. CRootBox can model fully grown root systems. Therefore, the model was enhanced to describe the emergence of basal and shoot borne roots. In this way the model is now capable of simulating the life span of dicotyledons and monocotyledons plants.
3. CRootBox enables root function modelling: The most important ways to couple root growth with soil properties are predefined. Soil properties can influence growth direction, root elongation rate, branching patterns, and the angle of lateral root emergence. The development of root functional models is simplified by the Python library of CRootbox which makes it easy to glue the different sub-models together.

### 4.1 CRootBox enables the modelling of mature root systems of a large range of plant species

Roots are important components of the global ecosystems. From a crop production perspective, they are responsible for the acquisition of water and nutrient and, as such, key to plant productivity. From an ecological perspective, roots play an important role for the soil water and carbon cycles, soil stability, the soil fauna, etc. Root models can help to better understand the quantitative role of roots in the ecosystem. It is therefore important for these models to be able to represent a wide range of root systems, without being limited to crop plants.

The modular structure of CRootBox enables the simulation of virtually any type of root system. For any root system type, only a limited number of input parameters are required, most of them being relatively easy to acquire experimentally (e.g. from excavation experiments). In the database we created, we provide 22 parameter sets for 14 different species based on published parameters. Those parameter sets of a wide variety of species are made easily accessible through the web application and a figshare collection, and we expect to update that collection to encompass more and more species.

The maximal rooting depth that can be reached by any root system is limited by the maximal root length of the main roots. Root parameters gained from images of young root systems may underestimate this important parameter; field scale simulations with virtual coring may help to achieve realistic root architecture parameters for mature plants.

The standard deviations of the different model parameters determine how different the individual realisations may be from each other. Image analysis results of rhizotron images suggest that we can expect a large standard deviation for root architectural parameters.

The root growth modelling in containers based on signed distance functions allows to mimic experiments that use specific containers, also split-root boxes, and it also works to simulate root growth around obstacles. It may for example be helpful to anticipate wall effects in a given container size.

CRootBox provides an interface to simulate any user-defined type of tropism or response to soil conditions locally experienced by the root system. This is demonstrated in Fig. 12; this example is based on the class *SoilPropertySDF* where a layer or a patch of elevated soil concentrations is defined via a signed distance function. Other possibilities such as passing the information on the soil property on a certain grid may be derived from the base class *SoilProperty*.

A limitation of the CRootBox may be seen in the fact that it does currently not explicitly compute during the simulation secondary root growth or variable root diameter along the branches. However, this can easily be computed a-posteriori, e.g. based on root segment age.

### 4.2 CRootBox enables root system modelling at the field scale

Roots do not grow alone in the soil. They grow within a plant community and influence (and are influenced) by their direct neighbours. They compete with each other for the same soil resources (water and nutrients) and can present complementary development strategies to maximise such resource acquisitions. To take this interaction into account, models should able to simulate several plants at the same time, within the same physical domain. Other root architecture models have been used to simulate a few plants simultaneously, e.g. SimRoot, R-SWMS, however, to our knowledge, no explicit root architecture model has so far been used at the field scale, simulating several hundred plants simultaneously.

CRootBox was entirely redeveloped in C++, an efficient programming language, to be able to simulate not only single root systems (as was previously the case with the Matlab version), but also field scale simulation, that encompass more than 200 individual mature root systems. So far, this kind of large scale modelling has only been used to simulate structure only together with related metrics like root length density distribution. In the future, fields scale modelling will allow us to investigate soil-root interactions from an ecological perspective. For instance, what is the functional importance of developmental plasticity within a single genotype, or how can we leverage complementarity when combining different crops species within the same field.

### 4.3 The object oriented structure of CRootBox will enable the extension to a whole plant model

Nowadays, only a few functional-structural plant models are able to represent both the root and the shoot as a single network (Lobet et al. 2014, Janott et al. 2011). However, such connection is needed to better understand the complex interplays and trade-offs that plants have to face during their growth and development. In particular, water and carbon flows are tightly intertwined and have a mutual strong influence. The object oriented structure of CRootBox allows for a direct extension from a root to whole plant model.

### 4.4 The CRootBox-Python binding enables an easy and straightforward communication with environmental models

Different strategies exist when it comes to combining plant and environmental simulations. The first strategy is to couple both within a single model. The second is to use separated models and couple them through a common interface.

For CRootBox, we decided to leverage the second strategy, to allow a greater modularity. Several environmental models exist, each with their own strengths and weaknesses and, depending on the research question at hand, one might want to use one or the other. With that in mind, we developed CRootBox as a single module, that could, in theory, be combined to any type of environmental model. We used Python as a glueing language between CRootBox and other models, as was done similarly in the past (Pradal *et al*. 2008). As demonstrated with soil physics with Python model, that strategy allows us to easily do the coupling.

### 4.5 Concluding remarks

In conclusion, we present a fast and flexible functional-structural root model which is based on state-of-the-art computational science methods. It is open source and available via a github repository. It is the first root architecture model which provides control of the segment length and hence spatial discretisation of the root architecture as numerical grid. Furthermore, it is the first root architecture able to simulate explicitly many root architectures on the field scale. CRootBox facilitates modelling of root responses to environmental conditions as well as root effects on soil. In the future, we plan to extend this approach to include mycorrhization (Schnepf *et al*. 2016) and to the above-ground.

## Appendix A Doussan model and its numerics

The derivation is based on Doussan et al., 2006 and Roose and Fowler, 2004.

### Model derivation

The axial water flux, q_z_, in the xylem of one root segment is given by

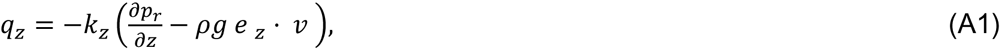

with units [L^3^ M^-1^ T^-1^], see Eqn 3.1, Roose and Fowler, 2004. The parameter *k_z_* is the axial conductance [L^5^ T], *p_r_* is the pressure inside the xylem [M L^-1^ T^-2^],*ρ*is the density of water [M L^-3^], *g* is the gravitational acceleration [L T^2^], *e_z_* the downward unit vector [1], and *v* [1] the normalised direction of the xylem. Thus Eqn (A1) can be expressed as

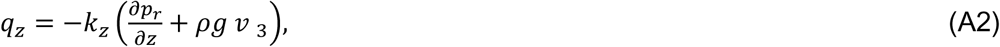

where *v*_3_ is the *z*-component of the normed xylem direction.

The radial flux is given by

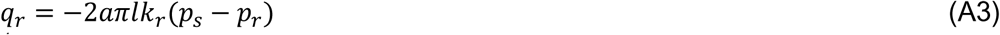

with units [L^3^ T^-1^] (based on Eqn 3.3, Roose and Fowler, 2004), where *a* is the root radius [L], *l* is the segment length [L], *k_r_* is the radial conductance [L^2^ T M^-1^], and *p_s_* is the pressure of the surrounding soil [M L^-1^ T^-2^].

Therefore, the net flux is

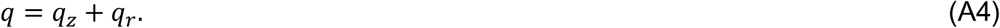

In a mathematical graph that represents the root system for each node *i* the sum of fluxes must be zero (first Kirchhoff’s law)

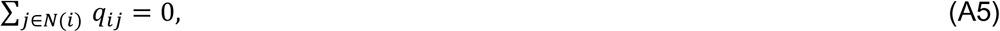

where *N*(*i*) are the nodes connected to node *i* and *q_ij_* is the net flux of the edge connecting node *i* and node *j*.

### Discretisation

In the graph the pressure *p_i_* is defined for each node *n_i_*. The edges at node *n_i_* are denoted as *e_ij_* with *j* ∈*N(i)*, where *N(i)* are the indices of the neighbouring nodes (the root collar and the root tips have one neighbour, and branch points have three neighbours). Thus, the edge *e_ij_* connects node *n_i_* and node *n_j_* for each *j* ∈*N(i)*.

For each edge *e_ij_* the axial water flux from *n_i_* to *n_j_* is

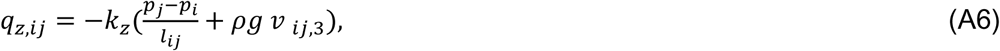

and the radial flux from segment *e_ij_* into the soil is

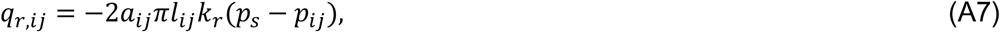

where *l_ij_* is the length, *v_ij_* the normed direction, *a_ij_* the radius, *p_ij_* is the mean edge pressure *p_ij_* = *0.5(p_i_+p_j_)* of the edge *e_ij_*. The value *p_s_* is the soil potential, surrounding the edge *e_ij_*. Therefore, the net flux of *e_ij_* is given by

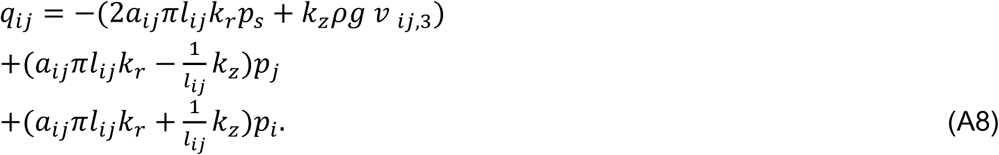

Eqn (A5) states that all fluxes into each node cancel out. This can be presented as a linear equation

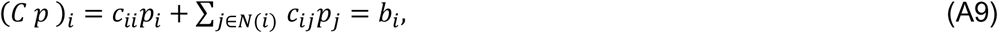

where each row *i* of matrix *C* represents the linear equation for node *i*. The diagonal elements of *C* are derived by the third line of Eqn (A8):

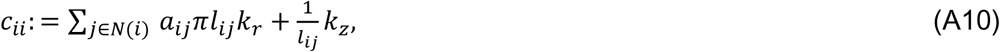

and the other entries by the second line of Eqn (A8):

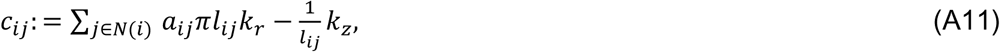

The value *b_i_* is derived from first line of Eqn (A8):

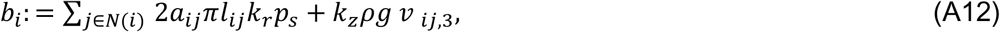

This yields the linear system *Cp*=*b* of Eqn (A9).

Note that *C* is symmetric (since the graph is undirected) and sparse (most *c_ij_* are zero, all which are not connected by an edge *e_ij_*). The soil matric potential *p_s_* and the direction of the edges *v_ij_* only enter the equation on the right hand side *b_i_*.

### Boundary conditions

For simplicity we assume a no-flux boundary condition at the root tips. This is a simplification, however, water can enter or leave radially in the edge representing the root tip. Therefore, the root tip conductivity can be easily adjusted by changing this edges root radial conductivity *k_r_*.

For this reason the only important boundary condition is at the root collar. Either a Dirichlet boundary condition (fixed potential) or Neumann boundary condition (fixed flux) is used.. Furthermore, often a combination is applied, where a potential flux is predetermined, but the boundary condition is switched to Dirichlet if the pressure magnitude becomes unreasonable high. In the following we assume the top node has index 1.

### Dirichlet

The simplest way to implement a fixed pressure at node 1, is to replace row 1 in the matrix *C* by *e*_1_^*T*^, and *b*_1_ by the desired matric potential *h_top_*. In this way the equation for node 1 of the linear equation *Cp*=*b* reads *p*_1_=*h_top_*.

### Neumann

Flux in node 1 is implemented by adding the flux to *b_1_*. Therefore, the net flux of row one does not sum up to 0 (see Eqn A5) but equals the desired flux.

## Appendix B Database for root system architectures

Table B1 presents an overview of literature sources which were used to parameterize the CRootBox model. The following paragraphs explain how we handled and substituted missing CRootBox parameter values.

**Table B1.**
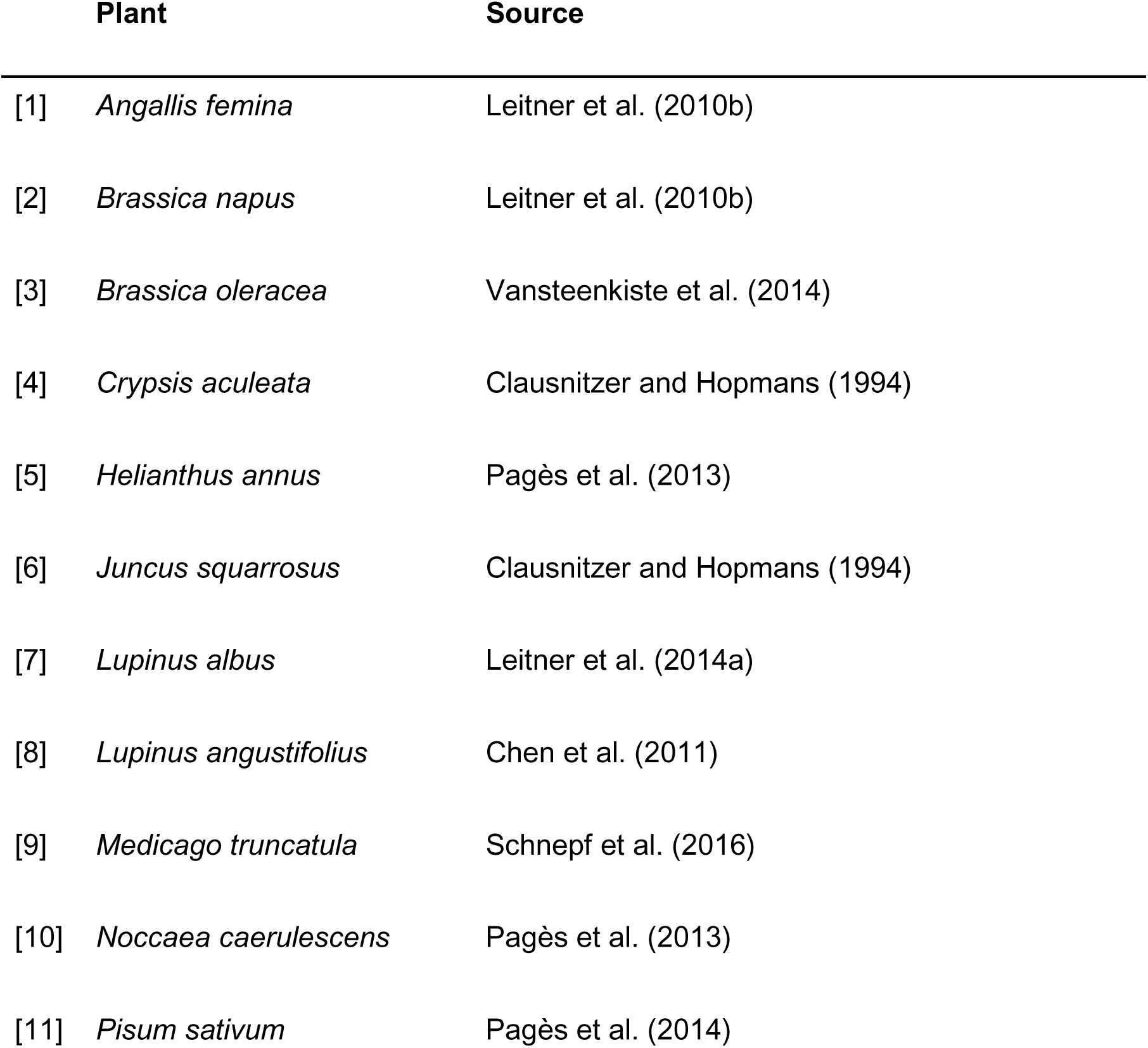
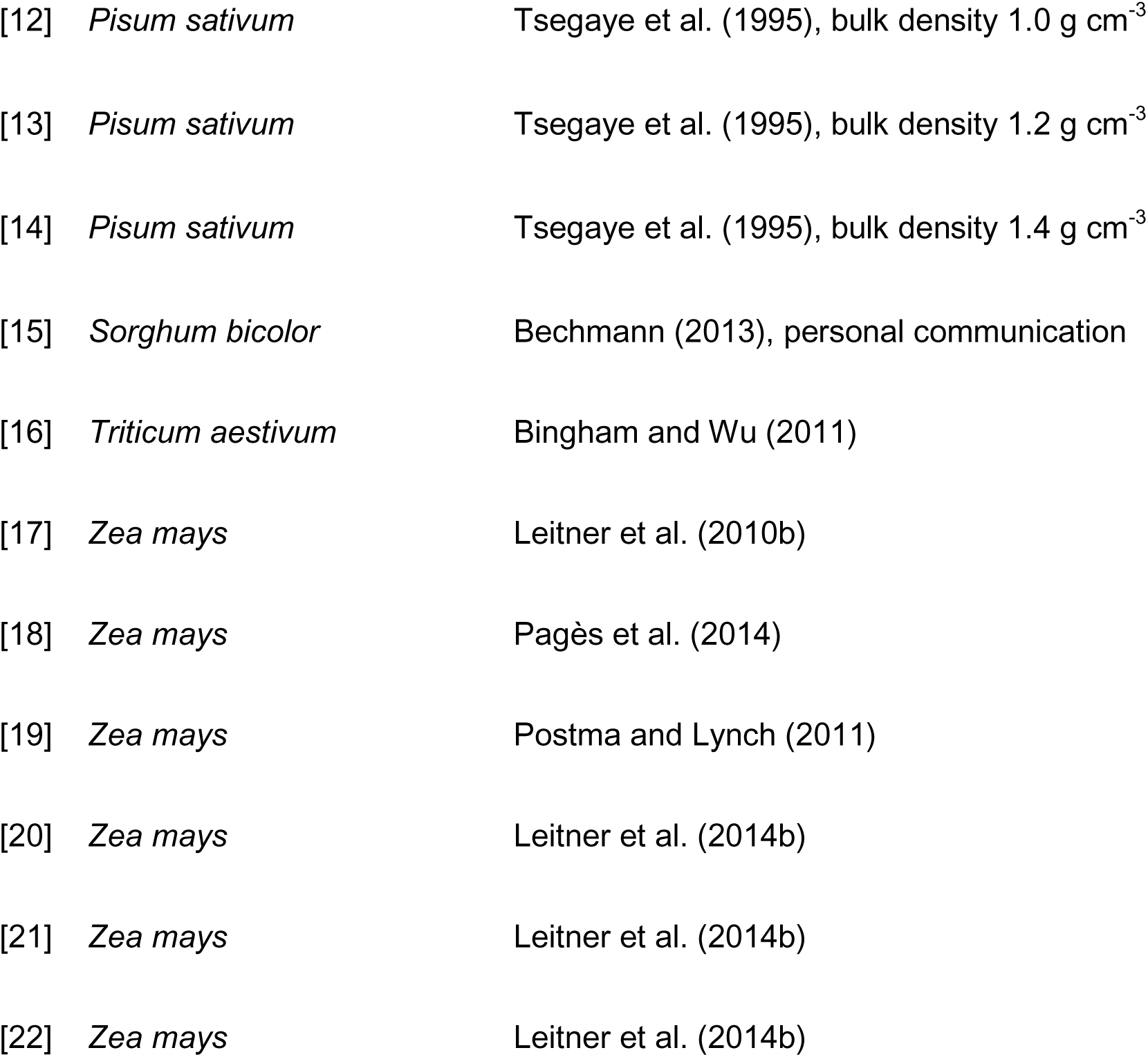
Literature sources used to parameterize the CRootBox model. Numbers in brackets correspond to the numbers indicated in Fig. 5 and 6.

### Anagallis femina

The root system parametrization for *A. femina* is presented in Leitner et al. (2010b). The root system was parametrized by visual comparison with images published by Kutschera (1960). All parameters for CRootBox are provided in the paper.

### Brassica napus

The parameters for *B. napus* were obtained by Leitner et al. (2010b) based on the drawings of Kutschera (1960). They observed two different kinds of lateral roots: near the surface the lateral roots are very dense and short and show plagiotropism, whereas the deeper lateral roots are long and show strong gravitropism. All parameters for CRootBox are provided in the paper. The parameters in the database are from Table 4 of Leitner et al. (2010) for *B. napus* (a).

### Brassica oleracea

Vansteenkiste et al. (2014) used the RootTyp model (Pagès et al., 2004) and evaluation of field experiments to determine root growth parameters. They considered three different root orders: main roots, long laterals from main roots, and short laterals from both main roots and long laterals. Missing parameters for CRootBox were the branching angles, and the length of apical and basal zones. We adjusted the unavailable parameters by visual comparison with other cauliflower root systems shown by Weaver and Bruner (1927) and Pagès et al. (2004).

### Crypsis aculeata

Root system parametrization presented in Clausnitzer and Hopmans (1994) was parametrized by visual comparison (Kutschera *et al.*, 1982). Missing parameters for the CRootBox model were the root radius, length of apical and basal zones, and the tropism parameters. They were substituted by visual comparison such that the newly simulated root system resembled that of the original publication.

### Helianthus annuus

Based on experimental data, Pagès *et al*. (2013) developed a stochastic 3D root system architecture model with the same general characteristics as presented in Pagès (2011). We used the parameters from their Table 3, option 3. Root radius in CRootBox is fixed for each root order; we used the average value of the minimal and maximal root radius given in Pagès et al. (2013), Table 3. The elongation was computed from their parameter “*E*” times the average root diameter. Apical and basal zone lengths needed by CRootBox were obtained by visual comparison such that the newly simulated root system resembled that of the original publication.

### Juncus squarrosus

Clausnitzer and Hopmans (1994) parametrized the root system of *J. squarrosus* by visual comparison with illustrations presented in Kutschera et al. (1982) considering two different root orders. Missing parameters for CRootBox were the root radius, length of apical and basal zones, and the tropism parameters. They were substituted by visual comparison such that the newly simulated root system resembled that of the original publication.

### Lupinus albus

Leitner et al. (2014a) parametrised 26 days old *L. albus* root systems from neutron radiography images. The parametrisation is not based on strict topological root orders but on root types that emerge with a certain probability. All parameters for CRootBox are provided in the paper.

### Lupinus angustifolius

Chen et al. (2011) measured root growth parameters for *L. angustifolius* by using semi-hydroponic bin systems. The authors did not perform simulations, but most parameters for CRootBox could be retrieved from these data. We substituted the missing values for maximal root length of first order laterals, length of apical and basal zones, elongation rate of first order laterals and the tropisms parameters with parameters for the plant *Lupinus albus* obtained by Leitner et al. (2014).

### Medicago truncatula

Schnepf et al. (2016) obtained CRootBox root architectural parameters for *M. truncatula* by direct measurement of images published by Bourion et al. (2014). All parameters for CRootBox are provided in the paper.

### Noccaea caerulescens

Based on experimental data, Pagès et al. (2013) developed a stochastic 3D root system architecture model with the same general characteristics as presented in Pagès (2011). We used the parameters from their Table 3, option 3. Root radius in CRootBox is fixed for each root order; we used the average value of the minimal and maximal root radius given in Pagès et al. (2013), Table 3. The elongation was computed from their parameter “E” times the average root diameter. Apical and basal zone lengths needed by CRootBox were obtained by visual comparison such that the newly simulated root system resembled that of the original publication.

### Pisum sativum

Four different parameterisations for P. sativum are in our database. The first parameter set stems from Pagès *et al*. (2014) for the model ArchiSimple. The second to fourth data sets stem from Tsegaye et al. (1995) for the model RootMap.

*P. sativum* [11] is derived from the parameters for the model ArchiSimple. No different parameterizations are given for the different root types; variations depend on the root radius. This makes it difficult to use this parameterisation for CRootBox; we retrieved the internodal distance, root radius, initial growth rate for the tap root from the paper, and substituted all other parameters by visual comparison such that the newly simulated root system resembled that of the original publication.

*P. sativum* [12]-[14] were derived for the model RootMap. The plants were grown under laboratory conditions in soil cylinders with three different bulk densities (1.0, 1.2, and 1.4 g cm^-3^). The parameters root length and root angle were measured after ten days and three root orders were specified. Missing values for CRootBox were maximal root lengths, apical and basal zone lengths, root radius and the tropism parameters. They were obtained by visual comparison such that the newly simulated root system resembled that of the original publication.

### Sorghum bicolor

This parameter set is based on parameters for the Root Typ model for *S. bicolor* (M. Bechmann, personal communication). Missing parameters for CRootBox were the length of apical and basal zones. We adjusted the unavailable parameters by visual comparison with other cauliflower root systems shown by Weaver & Bruner (1927) and Pagès et al. (2004).

### Triticum aestivum

This root system was parameterized based on Bingham and Wu (2011). Parameters missing for CRootBox were the maximal root length for the main roots, basal zone lengths, root radius, and the tropism parameters. They were estimated by visual comparison with the original publication.

### Zea mays

Six parameter sets for *Z. mays* are contained in our database.

*Z. mays* [17] parameterisation is based on Leitner et al. (2010a). All parameters for CRootBox are provided in the paper.

*Z. mays* [18] is parameterised based on Pagès et al. (2014) for the model ArchiSimple. No different parameterizations are given for the different root types; variations depend on the root radius. This makes it difficult to use this parameterisation for CRootBox; we retrieved the internodal distance, root radius, initial growth rate for the tap root from the paper, and substituted all other parameters by visual comparison such that the newly simulated root system resembled that of the original publication.

*Z. mays* [19] is parameterised based on Postma and Lynch (2011) for the model SimRoot. The parameters missing for CRootBox include apical and basal zone lengths, and the tropism parameters. We substituted them by parameters for the maize root system [18] from Pagès *et al*. (2014).

*Z. mays* [20–22] parameterisation is based on Leitner et al. (2014b), representing three different phenotypes of maize, one reference phenotype, one phenotype with steeper main roots, and one with steep main roots and with longer lateral roots. All parameters for CRootBox are provided in the paper.

## Supplementary data

S1: Doxygen documentation of the CRootBox code.

## Acknowledgements

This work was funded by the German Federal Ministry of Education and Research (BMBF) in the framework of the funding initiative “Soil as a Sustainable Resource for the Bioeconomy – BonaRes”, project “BonaRes (Module A): Sustainable Subsoil Management - Soil3; subproject 3” (grant 031B0026C) and by the German Research Foundation DFG (grant numbers PAK888, TR32-B4). C. Sheng has a PhD scholarship of the China Scholarship Council CSC.

